# Inhibition of release of intestinal extracellular vesicles in *Ascaris suum* and immune modulation by the anthelmintic ivermectin

**DOI:** 10.64898/2026.08.25.745816

**Authors:** Dongjie Liu, Paul D. E. Williams, Michael J. Kimber, Alan P. Robertson, Richard J. Martin

## Abstract

Ivermectin is an important broad-spectrum anthelmintic used to treat nematode parasites including gastro-intestinal infections of humans and animals. The mode of action for Ivermectin is understood to involve activation of inhibitory glutamate-gated chloride channels (GluCls). Ivermectin has also been reported to inhibit the release of extracellular vesicles (EVs). We found that EVs are released from the whole intestine of the gastro-intestinal parasite, *Ascaris suum*. Proteomic analysis identified 1,574 proteins within these intestinal EVs, including 96 nematode proteins with putative immune-associated functions based on homology to proteins involved in host immune processes and 130 proteins with predicted digestive functions. Comparative analysis following ivermectin exposure revealed 38 differentially abundant proteins that included the putative immune-related proteins: transthyretin-like proteins, a small heat-shock antigen, a phospholipase A2, and the NF-κB subunit p105. Thus, ivermectin modulated the potential immune-related cargo of intestinal EVs. The ivermectin inhibition of intestinal EV release was concentration-dependent with an *IC50* of 64 nM. We also identified the expression of GluCl subunit receptor genes in the *Ascaris* intestine. The potent inhibitory effect of ivermectin on the release of these EVs from the nematode intestine and the expression of GluCl channel subunits sheds further light on the site and mechanisms of action of this important anthelmintic.

**Graphical Abstract:** Extracellular vesicles have become recognized as important mediators of intercellular communication and host–pathogen interactions, but the tissue origins and control of helminth-derived EVs and the mechanisms regulating their release remain poorly understood and may include the E/S pore and anal pore. Ivermectin is an important drug for treating nematode parasite infections. Significantly, we find that release of intestinal EVs (*Int.)* that carry immune modulating protein cargo from the nematode parasite intestine to the host is inhibited by the anthelmintic ivermectin with glutamate gated chloride channels.

## 1. Introduction

Soil-transmitted helminths (STHs), including *Ascaris*, *Trichuris*, and hookworms, are among the most widespread human parasitic infections, with more than 1.5 billion people affected in low-income countries (WHO, 2023). According to recent global burden of disease estimates, STH infections continue to account for an annual loss of 1.38 million disability-adjusted life years (DALYs) (Chen et al., 2024). *Ascaris lumbricoides* is the most prevalent STH species, infecting an estimated 772–892 million people (CDC, 2024). These infections contribute to malnutrition, hinder physical and cognitive development, reduce learning in school children and the ability to work (WHO, 2023). In livestock, gastrointestinal parasites lead to significant economic costs, by reducing growth rates, poor feed conversion efficiency, exacerbating food insecurity and nutritional deficits in already impoverished communities (de Silva et al., 2003, Roepstorff et al., 2011, Thamsborg, 2013, Chen et al., 2024). Poor sanitation and the lack of effective vaccines means that prevention and treatment of STH infections relies on a drug from one of the three major classes of anthelmintics.

The macrocyclic lactones (ivermectin & avermectin) are one the major classes of anthelmintics and are used to treat nematode infections in both livestock and humans (Omura and Crump, 2004, Geary, 2005, Campbell, 2012). The major targets of macrocyclic lactones are the glutamate gated chloride channels (GluCls), which were functionally characterized from *Caenorhabditis elegans* (Cully et al., 1994, Arena et al., 1995). In *Ascaris suum*, GluCls have been identified in pharyngeal muscle and nerves and are sensitive to the macrocyclic lactones like ivermectin (Martin, 1996, Brownlee et al., 1997, Wolstenholme, 2012). The GluCl subunits *glc-2*, *glc-3*, *glc-4* & *avr-14* have since been identified in parasitic nematodes, including *B. malayi* (Lamassiaude et al., 2022).

In addition to the inhibition of nerves and muscles of the parasite, ivermectin also inhibits release by the whole worm of extracellular vesicles (EVs), which are implicated in host– parasite interactions and enhancement of parasite survival in the host (Moreno et al., 2010, Buck et al., 2014, Loghry et al., 2020). EVs are membrane-enclosed structures that contain proteins, small RNAs (miRNAs and circRNAs), lipids, and metabolites (EL Andaloussi et al., 2013, Minkler et al., 2022, Welsh et al., 2024). Helminth-derived EVs are proposed as key mediators of host–parasite interactions capable of modulating and suppressing host immune responses which facilitate long-term infection and survival (Marcilla et al., 2012, Buck et al., 2014, Tritten and Geary, 2018, Cortés et al., 2025). The source of these EVs and mechanism of release is not well understood. The ES pore located near the head of the nematode parasites has been suggested to be the major release site, although the intestinal epithelium and anal pore have also been proposed (Buck et al., 2014, Hansen et al., 2019, Loghry et al., 2020).

Helminth EV-associated miRNAs have been shown to induce Th2-type immune polarization in the host through pathways involving IL-10, IL-25, and IL-33, reducing pro-inflammatory responses (Locksley, 1994, Maizels and Yazdanbakhsh, 2003). In addition, circRNAs that interact with host miRNAs to regulate host gene expression at the host– parasite interface have been described (Memczak et al., 2013, Hansen et al., 2013, Minkler et al., 2022). Proteomic analyses of *A. suum* have revealed that these parasites secrete proteins that have potential roles in mediating host–parasite interactions (Chehayeb et al., 2014, Hansen et al., 2019). These proteins include heat shock proteins, proteases, kinases, oxidoreductases, peptidases, and glycolytic enzymes, indicating involvement in nutrient acquisition as well modulation of host immune responses (Chehayeb et al., 2014, Hansen et al., 2019). EVs also contain proteins linked to stress responses, proteolysis, and redox signaling as well as C-type lectins that may facilitate vesicle uptake by host cells (Hansen et al., 2019). These findings indicate that *A. suum* EV cargo is not restricted to host immune modulation but includes other housekeeping proteins. Release of the EVs from whole worms has been demonstrated to be inhibited by anthelmintic compounds, including ivermectin (Moreno et al., 2010, Harischandra et al., 2018, Loghry et al., 2020). The outcome of this anthelmintic inhibition is to reduce suppression of the host immune system by the parasite leading to parasite expulsion and death (Buck et al., 2014, Loghry et al., 2020). The locations and tissues for the inhibitory effects of ivermectin on the release of EVs remain to be determined and characterized.

In this study, we show that isolated intestines of *A. suum* secrete and are a source of EVs and that ivermectin inhibits EV release in a concentration-dependent manner with an *IC₅₀* of 64 nM. In addition, RT-PCR analysis demonstrated that the intestine expresses the GluCl subunit genes *Asu-glc-2, Asu-glc-3, Asu-glc-4,* and *Asu-avr-14*. Proteomic analysis showed that *A. suum* intestinal EVs contain over 1,500 proteins, including 96 putative immune-related proteins and 130 associated with predicted digestive functions. Following ivermectin treatment, our analysis revealed 38 differentially abundant proteins, some of which have potential immune modulatory functions. These results demonstrate the *Ascaris* parasite intestine is a major source of EV release involved in host immune suppression. We found that ivermectin inhibits the release of these EVs and alters their putative immune-related protein cargo.

## 2. Materials and Methods

### 2.1 Collection and maintenance of *A. suum*

Adult female *A. suum* worms were collected from the JBS Swift and Co. pork processing plant, Marshalltown, Iowa. Worms were maintained in *Ascaris* Ringers Solution (ARS: 13 mM NaCl, 9 mM CaCl₂, 7 mM MgCl₂, 12 mM C₄H₁₁NO₃, Tris, 99 mM NaC₂H₃O₂, 19 mM KCl, and 5 mM glucose pH 7.8) at 32°C for 24 h to allow for acclimatization before use in experiments. The worms were used the following day. All the worms were examined at the start of each experimental day and discarded if they were damaged or immotile.

### 2.2 Collection of Extracellular Vesicles (EVs) from *A. suum* Intestinal Sections

The whole intestine of adult *A. suum* was extracted from the parasite by dissecting the parasite from the base of the pharynx to the anal pore (Harpur, 1977). The intestine was removed from the body using fine forceps, avoiding other tissues including the reproductive organs and muscle bags. The intestines were either processed whole or dissected into three anatomical regions: 1) the anterior region, which is primarily involved in digestion stretching from the base of the pharynx to the vulvar pore; 2) the middle region which is associated with nutrient absorption and extends from the vulvar pore to the ovarian tissue; and 3) the posterior region from the ovarian tissue to the anal pore (Fig. 1A) (Harpur, 1977). The intestinal pieces were opened longitudinally, exposing the lumen, and thoroughly washed with *Ascaris* Perienteric Fluid (APF: 23 mM NaCl, 110 mM sodium acetate, 24 mM KCl, 1 mM CaCl₂, 5 mM MgCl₂, 5 mM HEPES, and 11 mM D-glucose; pH 7.6) to remove residual luminal contents and debris.

**Fig. 1:**
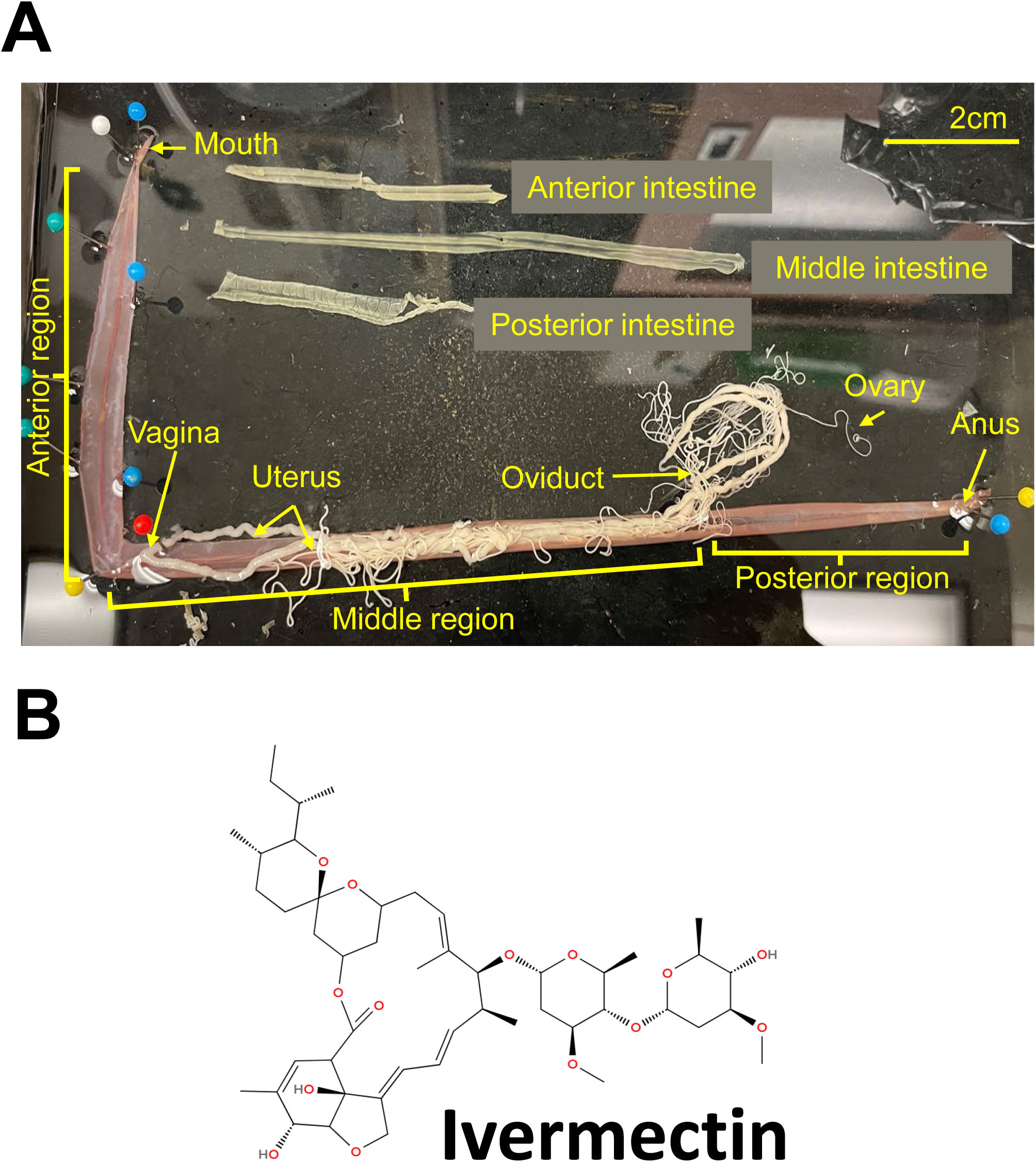
Dissected intestine of *Ascaris suum*. **A:** Photograph of dissected female *Ascaris suum* with the intestine separated from the main body and cut into the anterior, middle and posterior sections. Key structures of the anatomy are labelled. **B:** 2D skeletal structure of ivermectin

Intestinal sections were weighed and transferred into 15 mL tubes containing 10 mL APF supplemented with 0.01% dimethyl sulfoxide (DMSO) and were incubated at 37 °C for 4 hrs. to promote EV secretion. Following incubation, the intestinal sections were removed, and the conditioned APF solutions were collected for EV isolation and characterization. A total of fourteen independent worms as biological experiments were performed for all sections tested. For NTA analysis, conditioned APF solution from whole intestine, anterior, middle, and posterior intestinal sections were diluted 500-fold in filtered dPBS prior to particle quantification.

### 2.3 Measuring the Effects of Ivermectin on Intestinal EV Secretion

To assess the effect of ivermectin on EV secretion from the intestine, the anterior, middle, and posterior intestinal sections were divided equally into two halves (anterior and posterior), allowing for paired comparisons within the same worm. For half of the experiments, the anterior half of the intestinal section was incubated in 10 mL APF containing 0.01% DMSO, while the posterior half was incubated in 10 mL APF containing 1 µM ivermectin (Fig. 1B). In the remaining experiments, the treatment order was reversed, with the anterior half of the intestinal section being treated with 1 µM ivermectin and the posterior half treated in 0.01% DMSO. All samples were incubated at 37 °C for 4 hrs. A total of fourteen independent biological experiments were performed for all sections tested. For NTA analysis, conditioned APF solutions from ivermectin and DMSO treated anterior, middle, and posterior intestinal sections were diluted 100-fold in filtered dPBS prior to particle quantification.

For dose-response experiments the anterior sections of *A. suum* intestines were divided into six 1 cm pieces. For all experiments, samples were incubated in either 10 μM, 1 μM, 100 nM, 10 nM, 1 nM, and 100 pM ivermectin, at 37°C for 4 hrs. For three experiments, the samples were treated with the piece nearer the mouth being exposed to the highest ivermectin concentration and the piece near the vulvar pore being treated with the lowest concentration. To control positional bias in the remaining three experiments, the assignment was reversed with the piece near the vulvar pore being exposed to the highest concentration of ivermectin and the piece near the mouth being exposed to the lowest. For control experiments the anterior intestines of six individual *A. suum* were dissected into six 1 cm pieces and incubated in APF containing 0.01% DMSO at 37 °C for 4 hrs. At the end of incubation, all solutions were collected for EV isolation and quantification. For NTA analysis in the ivermectin dose-response experiments, conditioned APF solutions from anterior intestinal sections were diluted 100-fold in filtered dPBS prior to particle quantification.

### 2.4 EV isolation, quantification and characterization

After incubation, conditioned solutions were passed through a 0.2 µm PVDF syringe filter (GE Healthcare, Chicago, IL) and ultracentrifuged at 120,000 × g for 90 mins at 4°C. Supernatants were decanted, leaving approximately 1.5 mL above the pellet to avoid disturbance. The EVs contained in the pellet were purified by size-exclusion chromatography (SmartSEC™ single columns), and the EV-containing fractions were concentrated by ultracentrifugation at 186,000 × g for 2 hrs. at 4°C. The pellets were resuspended in 500 µL of sterile-filtered dPBS (Thermo Fisher Scientific). Samples were diluted in filtered dPBS to achieve particle counts (20-120 particles per frame). Dilution factors were optimized for each experiment and are indicated in the corresponding sections, 2.2 and 2.3. For each sample, videos of 5 x 60 sec were recorded at a camera level of 12, and particles were detected using a detection threshold of five._The EV concentration and size distributions were determined by nanoparticle tracking analysis (NTA; Nano-Sight LM10, Malvern Instruments, Malvern, UK)

### 2.5 EV visualization by Transmission Electron Microscopy (TEM)

Aliquots of purified EVs were fixed in 3% glutaraldehyde + 1% paraformaldehyde in 0.1 M cacodylate buffer. 200-mesh copper grids with carbon film (EMS) were glow-discharged (Pelco easiGlow, Ted Pella) to render a hydrophilic surface. 2 µL of fixed EV suspension were applied to each grid for 30s, excess wicked with filter paper, and immediately negative-stained with 2% (w/v) uranyl acetate (2 µL, 30s), then wicked and air-dried. Grids were examined on a JEOL 2100 transmission electron microscope operated at 200 kV and imaged with a Gatan OneView camera at the Iowa State University Light and Electron Microscopy Facility.

### 2.6 Proteomic Analysis of EV Contents

EV protein lysates were prepared from EV pellets prior to LC–MS/MS proteomic analysis. EV pellets were kept on ice and resuspended in 30 µL of 8 M urea prepared in 50 mM ammonium bicarbonate buffer (ABC, pH 8.5). The urea–ABC lysis buffer was prepared at room temperature without heating and deionized using AG 501-X8 resin before use to reduce ionic contaminants and minimize urea-derived carbamylation. EV pellets were lysed by repeated pipetting, vortexing, and incubation at room temperature. Specifically, each sample was pipetted up and down approximately 100 times, vortexed for 1 min, incubated at room temperature for 10 min, and vortexed again for 30 s. This cycle was repeated twice to promote complete solubilization of EV-associated proteins. The lysates were then centrifuged at 12,000–16,000 × g for 10 min at 4 °C to remove insoluble debris. The resulting supernatant was collected as the soluble EV protein lysate for downstream proteomic analysis. When protein quantification was required, aliquots were diluted to reduce the urea concentration before measurement or analyzed using a urea-compatible protein assay (Wiśniewski et al., 2009, Rontogianni et al., 2019). Samples were maintained on ice or at 4 °C for short-term handling and were stored at −80 °C if same-day processing was not possible, with repeated freeze–thaw cycles avoided.

### 2.7 Protein digestion and LC–MS/MS analysis

Crude EV protein extracts were processed for bottom-up proteomic analysis by in-solution enzymatic digestion. Briefly, protein extracts were reduced with dithiothreitol (DTT), and cysteine residues were alkylated with iodoacetamide prior to enzymatic digestion. Samples were then digested overnight with trypsin/Lys-C. Digestion was stopped by the addition of formic acid, and the samples were dried using a SpeedVac concentrator. Peptides were desalted using C18 columns (BioPureSPN Plate, HHNFR S18V; Nest Group) and dried again in a SpeedVac. Peptide concentration was determined using a bicinchoninic acid assay kit (BCA-1; Sigma-Aldrich). Peptide Retention Time Calibration standard mixture (PRTC; Pierce, part No. 88320) was spiked into each sample as an internal control for LC–MS/MS performance monitoring. Samples were normalized to 200 ng/µL peptide and 50 fmol/µL PRTC, and 2 µL of each sample was injected for LC– MS/MS analysis.

Peptides were separated by liquid chromatography using a Thermo Scientific Vanquish Neo UHPLC system and analyzed by tandem mass spectrometry on a Thermo Scientific Orbitrap Astral mass spectrometer equipped with a Thermo Scientific EASY-Spray ion source and column. The resulting intact and fragmentation pattern is compared to a theoretical fragmentation pattern using CHIMERYS. CHIMERYS was used to identify and quantify peptides from complex tandem mass spectra, including spectra containing co-fragmented peptide ions. Protein identification and quantification were performed based on peptide-spectrum matching and comparison of the observed MS/MS fragmentation patterns with theoretical fragmentation patterns generated from the protein database. The complete proteomic dataset, including protein identifications, peptide counts, and grouped abundance values for all DMSO and ivermectin treated replicates, is provided in Supplementary Data File S1.

### 2.8 Functional classification of extracellular vesicle proteins

To characterize the functional composition of the intestinal extracellular vesicle proteome, identified *Ascaris suum* proteins were assigned to functional categories using a priority-based scheme that combined UniProtKB (release 2026_02; accessed June 2026) annotation with manual curation. For each protein, the UniProtKB protein name and Gene Ontology (GO) biological process terms were retrieved and grouped according to their GO biological process terms. All remaining proteins lacking GO annotation were classified separately. The curated keyword lists were compiled from protein families previously reported in nematode and helminth secretome and extracellular vesicle studies, and all family assignments were manually reviewed. *A. suum* is not represented in standard GO-slim enrichment databases (e.g., PANTHER), so a curated approach was applied in place of automated analysis. Category and sub-family counts were visualized as pie and donut charts.

### 2.9 Measurement of differential abundance analysis and visualization

Extracellular vesicle proteins were performed using the grouped protein abundances generated by Proteome Discoverer, comprising five ivermectin-treated and five DMSO-treated (vehicle control) replicates, with all abundances normalized to the PRTC spike-in standard. ’Master Protein’ (Any UniProt-mapped *A. suum*) were retained, and contaminant entries (Cont_) and PRTC peptides were removed. All *Ascaris suum* proteins were mapped to UniProt identifiers. For each protein, we measured the abundance values in each treatment group, and the valid values were checked. Proteins below two valid abundance values in either treatment group (only present in 1 worm) were excluded from the differential-abundance test as a reliable estimation was not possible. The grouped protein abundances from five ivermectin-treated and five DMSO-treated replicates were log₂-transformed. For each protein, the log₂ fold change was calculated as the difference between the mean log₂ abundance of the ivermectin groups and that of the DMSO groups (mean log2 DMSO, mean log2 IVM; log2 FC = mean log2 IVM - mean log2 DMSO). To determine significance the five IVM log2 vs five DMSO log2 were assessed using Welch’s *t*-test (unequal variance), and *p*-values were adjusted for multiple comparisons using the Benjamini–Hochberg false discovery rate (FDR). Proteins with *p* < 0.05 and log₂ FC > 1 were considered candidate differentially abundant proteins. For heatmap visualization, abundance values were converted to *z*-scores across the ten samples for each protein [z = (log2 value - mean)/SD, across the 10 samples (DMSO & IVM)]. Volcano plots were generated by plotting each protein log2 FC (x) vs -log10(p) (y).

### 2.10 *A. suum* cDNA synthesis and RT-PCR detection of Glutamate-gated Chloride (GluCl) channels

A 3 cm segment of the anterior body of adult female *A. suum* was excised and opened longitudinally. The intestine was separated from the body wall using two fine forceps and each tissue was transferred to 1.5 mL RNase-free tubes. Tissues were snap-frozen in liquid nitrogen and stored at −80°C until processing. RNA was isolated from frozen tissues that were homogenized separately in 1 mL TRIzol™ Reagent (Life Technologies, USA) using a mortar and pestle according to the manufacturer’s protocol. cDNA was synthesized from one microgram (1 μg) RNA per sample using SuperScript IV VILO™ Master Mix (Life Technologies, USA). Paired intestine and body wall tissues were collected from five individual adult female worms for RT-PCR analysis.

RT-PCR was performed to detect *Asu-glc-2*, *Asu-glc-3*, *Asu-glc-4*, and *Asu-avr-14* (Supplementary Table 1) in the cDNA pool of the intestine and body wall using primers targeting the coding region of each gene (Supplementary Table 2). The reference gene *Asu-gapdh* served as the positive control. Negative controls included enzyme, water, and both forward and reverse primers for the target gene with no cDNA template. RT-PCR detection was performed on cDNA derived from all five individual worms described above. The cycling conditions for PCR were an initial denaturation for 2 min at 95°C, followed by 35 cycles at 95°C for 30s, 60°C for 35s, 72°C for 45s, followed by a final extension period at 72°C for 10 mins using GoTaq® G2 Hot Start Green Master Mix (Promega, USA). The PCR products of each gene were then separated on an individual 2% Agarose gels containing SYBR® Safe DNA Gel Stain (ThermoFisher Scientific), for 60 mins at 100 V, followed by visualization under UV light to confirm the presence of the genes. All photographs were acquired using Visionworks™ software (Analytik Jena) with an exposure setting of 3s per 1 frame.

RT-PCR was performed to generate full length transcripts of *Asu-avr-14* from the cDNA of the parasite intestine using primers that targeted the start and end codon (Supplementary Table 1). The reference gene *Asu-gapdh* served as the positive control. Negative controls included enzyme, water, and both forward and reverse primers for *Asu-avr-14* with no cDNA template. Amplification was performed with Platinum SuperFi™ II Green PCR Master Mix (Thermo Fisher Scientific) under the following conditions: 98 °C 30s; 35 cycles at 98 °C 30 s, 59 °C for 20 s, 72 °C for 40 s and a final extension at 72 °C for 10 mins. Products were identified on a 1% agarose gel containing SYBR® Safe (Thermo Fisher Scientific) at 100 V for 60 mins, visualized under UV illumination, and imaged with VisionWorks™ (Analytik Jena; 3 s exposure). Target bands were excised and purified with the NucleoSpin Gel and PCR Clean-up kit (Macherey-Nagel) per the manufacturer’s instructions. Purified amplicons were submitted to the Iowa State University DNA Sequencing Facility for Sanger sequencing (bidirectional reads were achieved using the same forward and reverse primers).

### 2.11 Statistical analysis

Statistical analyses were performed using Prism version 10.0 (GraphPad Software, La Jolla, CA, USA). EV concentrations were normalized to tissue wet weight (mg) for analysis. Differences in EV secretion among intestinal regions were analyzed using one-way ANOVA followed by a post hoc Šídák multiple-comparisons test. The effect of ivermectin on regional EV release was analyzed using an unpaired t-test, with a *P* value < 0.05 being considered statistically significant for all analyses. Dose–response relationships were analyzed using nonlinear regression with a log (inhibitor) versus response–variable slope model, and ivermectin concentrations were log10-transformed prior to analysis. A two-way ANOVA model was used to compare the means of EV size profiles following drug treatment, and statistical significance was determined using a post hoc Šídák multiple-comparisons test (*P* < 0.05 was considered significant). To account for the wide dynamic range of EV concentration values (particles mL^−1^ mg^−1^) across size subsets, a constant value of 1 was added to all EV concentration values (8 size subsets x 14 DMSO and ivermectin treated replicates per region) prior to log transformation, to avoid undefined values arising from zero counts in the log transformation. The adjusted values were them log10-transformed. A two-way ANOVA model was applied to the log10-transformed data to compare EV size profiles following drug treatment, and statistical significance was determined using a post hoc Šídák multiple-comparisons test (*P* < 0.05 was considered significant).

Proteomic data processing and statistical analyses were performed in Python using the pandas, NumPy, and SciPy packages, and figures were generated in R (version 4.5.2). Differential protein abundance between ivermectin and DMSO treated intestinal EVs was assessed by Welch’s *t*-test on log₂-transformed abundances to stabilize the variance, approximate a normal distribution and not assume equal variances between groups (n = 5 replicates). *P* values were corrected for multiple testing using the Benjamini–Hochberg false-discovery-rate (FDR) procedure. Proteins with *P* < 0.05 and log₂ fold change >1 were considered differentially abundant. Given the sample size, these proteins are reported as candidate differentially abundant proteins for functional interpretation. To ensure reproducibility, experiments were repeated independently, and details including the number of adult female worms used, intestinal preparations, drug concentrations, and treatment durations (IVM and DMSO) are provided in the corresponding figure legends.

## 3. Results

### 3.1 Characterization of extracellular vesicles from different regions of the intestine of *Ascaris suum*

The main source of EV release has been hypothesized to be the excretory/secretory (ES) pore that is near the head of the parasitic nematode. However, EV release may also occur from the anal pore by being released from the parasite’s intestine. To determine if the nematode intestines of the large pig gastrointestinal roundworm *A. suum* (Fig. 1A) release EVs, we dissected and isolated the intestine of the parasite. To collect the EVs from the dissected intestine we incubated them in 0.01% DMSO *Ascaris* Perienteric fluid (APF) for 4 hours at 37 °C. We obtained EV like particles with an average concentration/wet weight of 4.52 x 10^9^ ± 2.5×10^8^ particles mL^−1^ mg^−1^ (Fig. 2A; white bar; N=14). This suggested that the intestine is an important source of released EVs.

**Fig. 2.**
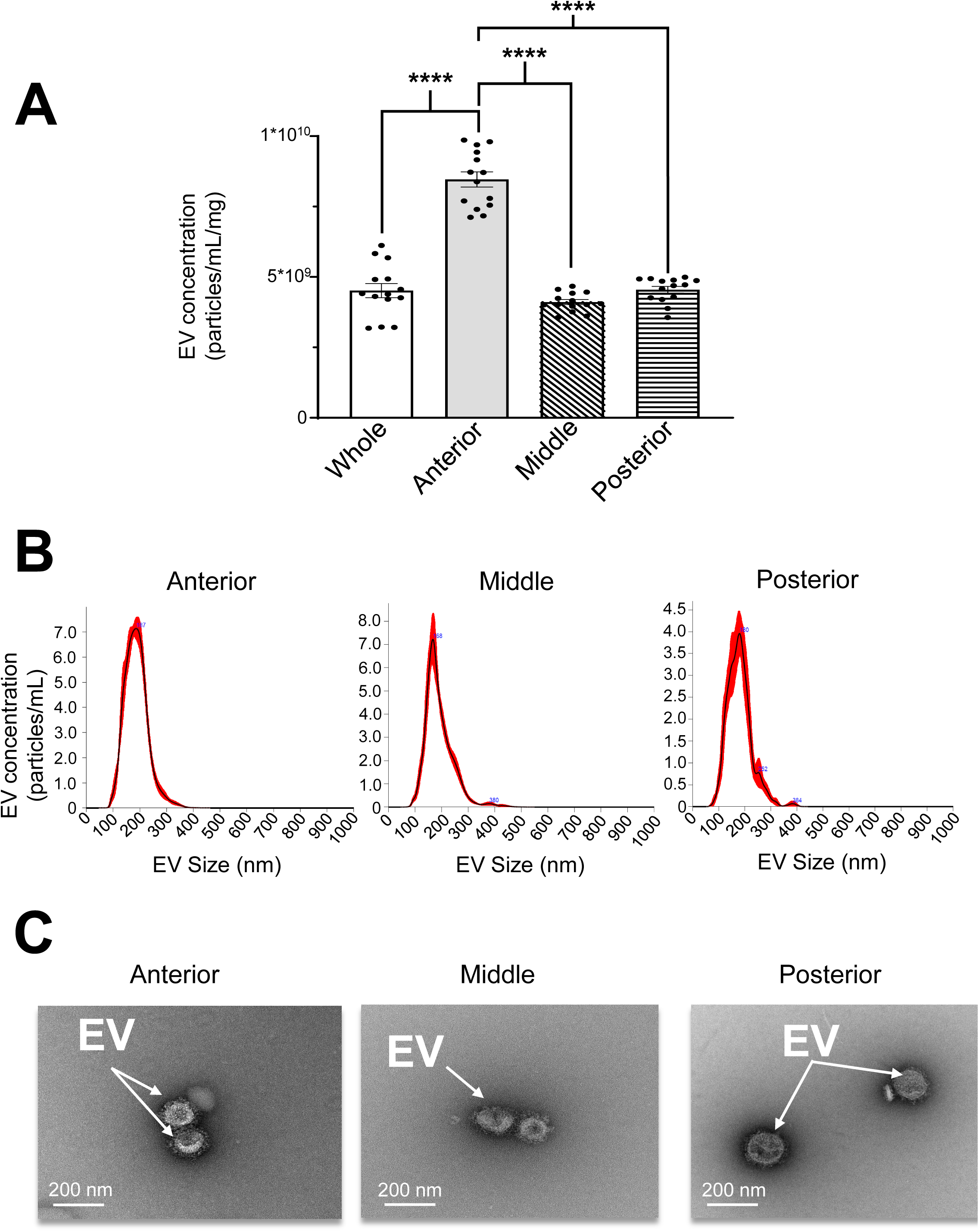
Regional concentrations of intestinal extracellular vesicles (EVs) from *Ascaris suum*. A: Quantification of EV concentrations released from the whole intestine and from anterior, middle, and posterior intestine regions. Analysis revealed the mean ± SEM concentrations/wet weight of 4.52 x 10^9^ ± 2.5×10^8^ particles mL^−1^ mg^−1^ from whole intestine, 8.46 x 10^9^ ± 2.7 ×10^8^ particles mL^−1^ mg^−1^ from anterior intestine, 4.11 x 10^9^ ± 9.0 ×10^7^ particles mL^−1^ mg^−1^ from middle intestine, and 4.55 x 10^9^ ± 1.18 ×10^8^ particles mL^−1^ mg^−1^ from posterior intestine. Statistical significance is indicated by **** for *P* < 0.0001. EV concentrations per mg in the anterior region were significantly higher than those in the whole intestine, middle intestine, and posterior intestine (P < 0.0001). means ± SEM, *N* = 14. **B:** Finite track length adjustment (FTLA) concentration/size distribution graphs from nanoparticle tracking analysis (NTA) of EV-like particles derived from the anterior (left), middle (center), and posterior (right) intestinal regions. Each graph represents the average FTLA profile. **C:** Transmission electron microscopy (TEM) images showing EVs (100–200 nm) released into culture media by the anterior, middle, and posterior regions of the intestine from adult female *A. suum* after 4 h incubation in *Ascaris* perienteric fluid containing 0.01% DMSO (pH 7.6). EVs were isolated and purified using ultracentrifugation, SEC columns, and 0.2 µm filtration. Scale bars are indicated. Original uncropped TEM images are presented in supplementary Fig. S2.

We sought to determine if the release of the EVs was evenly distributed along the entire length of the intestine or if the release was limited to a specific region. We separated the intestine of *A. suum* into distinct regions that were weighed wet. We divided the intestine into: 1) the anterior region; 2) the middle region; and 3) the posterior region. We again incubated each region in 0.01% DMSO APF for 4 hours at 37 °C and detected EV like particles from all three regions of the intestine (Fig. 2A). The anterior region had the highest concentration/wet weight (8.46 x 10^9^ ± 2.7 ×10^8^ particles mL^−1^ mg^−1^; grey bar; N=14), with the middle (4.11 x 10^9^ ± 9.0 ×10^7^ particles mL^−1^ mg^−1^; slanted bar; N=14) and posterior regions having similar concentrations (4.55 x 10^9^ ± 1.18 ×10^8^ particles mL^−1^ mg^− 1^; dashed bar; N=14). The EV sizes ranged between 50-400 nm (Fig. 2B). The anterior region model size was 187 nm, the middle region model size was 168 nm, and the posterior regions model size was 180 nm. The middle and posterior regions both had small auxiliary peaks near 380 nm with the posterior region also having a second auxiliary peak at 252 nm. The presence of auxiliary peaks suggests the presence of heterogenous subpopulations of EVs. We used Transmission Electron Microscopy (TEM) to observe the particles. We identified the presence of EVs based on the typical characteristic of a cup-shaped morphology (Fig. 2C and Supplementary Fig. S1) released from the anterior, middle, and posterior regions of the intestines.

### 3.2 Ivermectin inhibits intestinal EV secretion in *Ascaris suum*

Ivermectin has been found to inhibit release of EVs from whole nematode parasites [*Brugia malayi* from the ES pore, (Moreno et al., 2010); whole *A. suum,* (Loghry et al., 2020)] but it was not known if ivermectin inhibits the release of EVs from the intestine.

Initially we tested the effects of ivermectin on whole *A. suum* and confirmed that 1 µM ivermectin significantly reduces the EV concentration compared to untreated worms over 24 hours (Supplementary Fig. S2).

To determine if ivermectin suppresses the release of EVs from the intestine, we dissected again separated the intestine into the anterior, middle, and posterior regions. We then separated each region into two halves, with one half being incubated in APF with 0.01% DMSO for 4 hours at 37 °C, while the other half was incubated in APF containing 1µM ivermectin with 0.01% DMSO. We observed that 1 µM ivermectin significantly reduced the concentration of EVs per mg in all three regions of the intestine compared to untreated counterparts (Fig. 3 A).

**Fig. 3.**
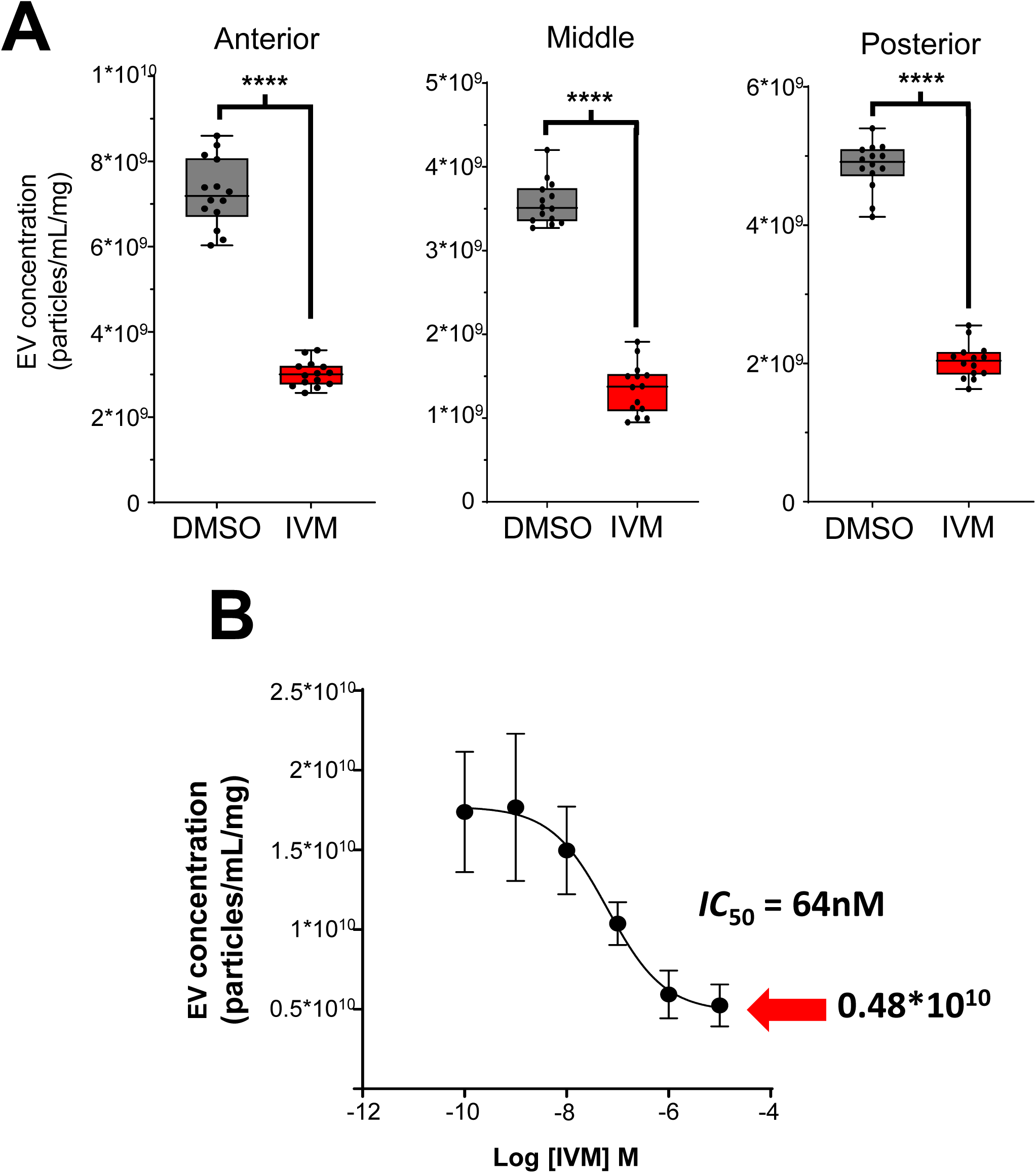
Ivermectin inhibits extracellular vesicle (EV) secretion from the intestine of adult female *Ascaris suum*. **A:** Adult female intestinal tissues (anterior, middle, and posterior regions) were cultured at 37 °C in *Ascaris* perienteric fluid with 1 µM ivermectin (IVM) or 0.01% DMSO. Media was collected after 4 h, and EVs were isolated and quantified by nanoparticle tracking analysis. EV concentrations obtained from the three intestinal regions of 14 independent worms, normalized to their corresponding intestinal wet weight. *Anterior intestine:* EV concentrations: Means ± SEM 7.26 x 10^9^ ± 2.15×10^8^ particles mL^−1^ mg^−1^ for control and 3.02 x 10^9^ ± 8.03×10^7^ particles mL^−1^ mg^−1^ for 1 µM ivermectin (IVM). **** *P* < 0.0001, N = 14, unpaired *t-test*. *Middle intestine:* EV concentrations: Means ± SEM 3.57 x 10^9^ ± 6.98×10^7^ particles mL^−1^ mg^1^ for control and 1.35 x 10^9^ ± 8.06 ×10^7^ particles mL^−1^ mg^−1^ for 1 µM ivermectin. **** *P* < 0.0001, N = 14, unpaired *t-test*. *Posterior intestine:* EV concentrations: Means ± SEM were 4.85 x 10^9^ ± 9.25 ×10^7^ particles mL^−1^ mg^−1^ for control with 0.01% DMSO and 2.03 x 10^9^ ± 6.83 ×10^7^ particles mL^−1^ mg^−1^ for 1 µM ivermectin. \*\*\*\**P* < 0.0001, N = 14, unpaired *t-test*. **B:** Ivermectin dose-response for the anterior region of the intestine. The anterior region was dissected into six sequential 1 cm pieces from <u>pharynx to vulvar pore</u>, and each piece was incubated with an ivermectin concentration that ranged from 10 µM to 100 pM (10^−5^–10^−10^ M) for 4 h. In three experiments, tissues were treated from pharynx to <u>vulvar pore</u>, and in three experiments from <u>vulvar pore</u> to pharynx. EV concentrations obtained from anterior region of 6 independent worms were normalized to the corresponding intestinal wet weight. Ivermectin inhibited EV release in a concentration-dependent manner, with an estimated *IC₅₀* of 64 nM, N=6.

To determine how effective ivermectin is at inhibiting EV release, we performed a dose– response analysis on the anterior region of the intestine because this region released the highest concentration of EVs. We dissected the anterior intestine into six sequential 1 cm segments extending from the mouth to the vulvar pore, and each segment was incubated for 4 hours with a different concentration of ivermectin ranging from 100 pM to 10 µM. EV secretion was quantified as before and normalized to tissue wet weight. Ivermectin inhibited EV release in a concentration-dependent manner, with an *IC50* of 64 nM (Fig. 3B; N=6). These findings demonstrate the intestinal EV secretory machinery is potently inhibited by ivermectin. Ivermectin did not inhibit release of all the EVs, leaving 4.8*10^9^ particles mL^−1^ mg^−1^ in 10µM ivermectin, and reducing the concentration from 1.77 *10^10^ particles mL^−1^ mg^−1^ in 100pM ivermectin (Fig. 3B). Thus, 27% of EVs from the intestine were not inhibited by ivermectin, suggesting the presence of an ivermectin-insensitive EV release pathway alongside the greater GluCl-sensitive pathway.

### 3.3 Ivermectin inhibits, but not equally, secretion of EV size subsets

It was of interest to determine if the inhibitory effect of ivermectin varied with the EV sizes or if all EVs were equally inhibited. Treatment with 1 µM ivermectin showed an inhibitory effect on EV secretion across multiple size ranges and from each of the three intestinal regions of *A. suum* (Fig. 4).

**Fig. 4.**
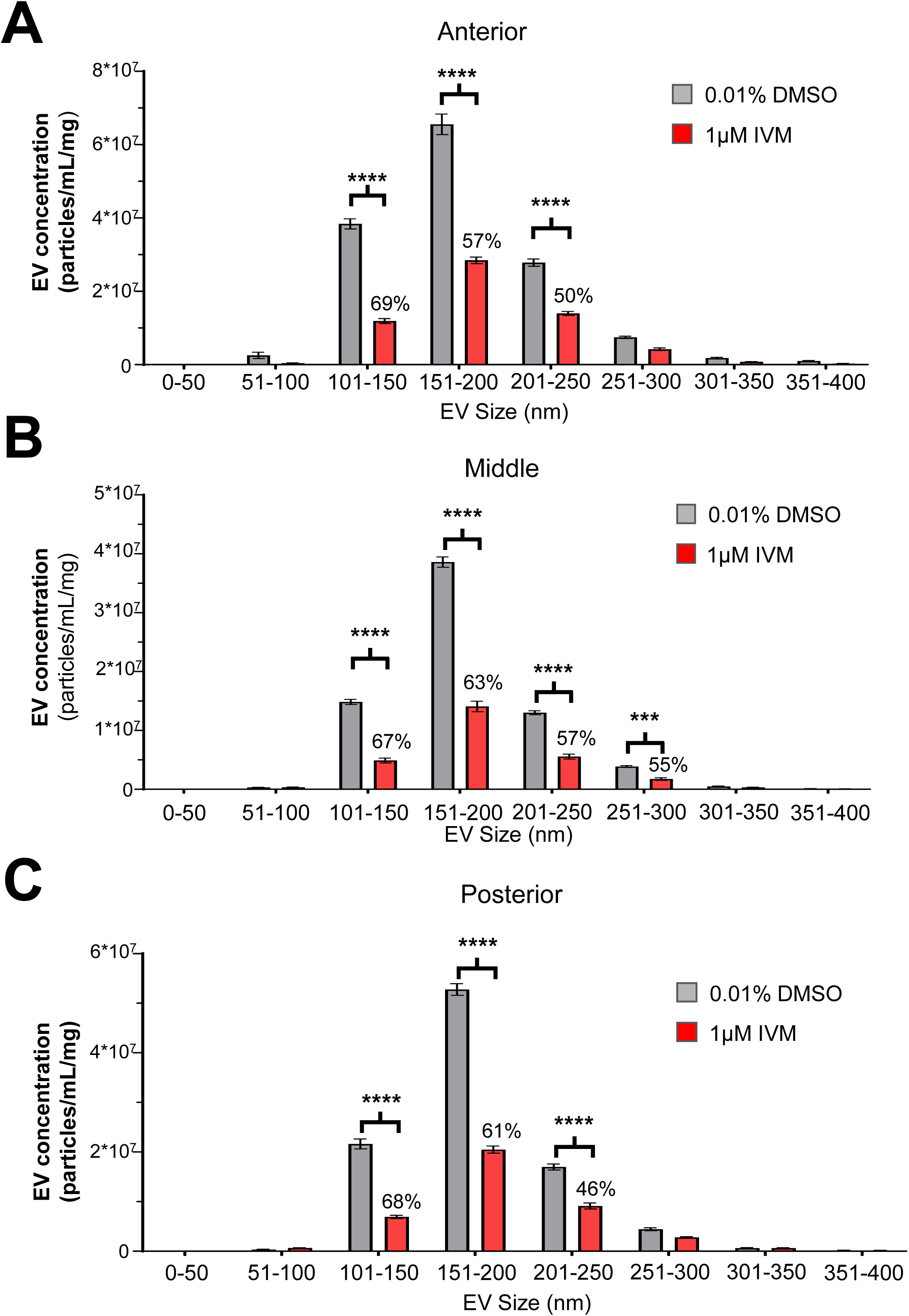
Secretion of distinct EV size subsets are inhibited by ivermectin. Adult female intestinal tissues (anterior, middle, and posterior regions) were cultured at 37 °C in *Ascaris* perienteric fluid with 1 µM ivermectin (IVM) or 0.01% DMSO (pH 7.6). Media was collected after 4 h, and EVs were isolated and quantified. A two-way ANOVA model was used to compare the means of EV size profiles following drug treatment, and statistical significance was determined using a post-hoc Šídák multiple-comparisons test (*P* < 0.05 was considered significant, *** for *P* <0.001, **** for *P* <0.0001). Data are presented as mean ± SEM, *N* = 14. Comparisons without significance markers were not statistically significant. **A:** In the anterior intestine, ivermectin (1 µM) significantly inhibited the 101–150 nm, 151–200 nm and 201–250 nm EV subsets. EV concentrations were 3.83 x 10^7^ ± 1.38 ×10^6^ particles mL^−1^ mg^−1^ with 0.01%DMSO and 1.19 x 10^7^ ± 6.78 ×10^5^ particles mL^−1^ mg^−1^ with 1 µM ivermectin in the 101-150 nm EV subsets; EV concentrations were 6.55 x 10^7^ ± 2.81 ×10^6^ particles mL^−1^ mg^−1^ with 0.01%DMSO and 2.84 x 10^7^ ± 8.92 ×10^5^ particles mL^−1^ mg^−1^ with 1 µM ivermectin in the 151–200 nm EV subsets; EV concentrations were 2.78 x 10^7^ ± 9.78 ×10^5^ particles mL^−1^ mg^−1^ with 0.01%DMSO and 1.40 x 10^7^ ± 5.35 ×10^5^ particles mL^−1^ mg^−1^ with 1 µM ivermectin in the 201–250 nm EV subsets. Percent reductions in EV concentration (particles mL^−1^ mg ^−1^) following ivermectin treatment are indicated above the red bars. Ivermectin reduced EV concentration (particles mL^−1^ mg ^−1^) by 69% in the 101–150 nm EV size range, 57% in the 151–200 nm range, and 50% in the 201–250 nm range. Two-way ANOVA showed a significant inhibitory effect of ivermectin on EV concentration (P< 0.0001; F=136.45; DFn=1, DFd=208), and a significant interaction between the effect of ivermectin and EV size (P=0.0013; F=3.53; DFn=7, DFd=208), indicating that ivermectin did not reduce all EV size populations equally and had different quantitative effects across EV size groups, consistent with heterogeneity within the intestinal EV population. Post-hoc Šídák multiple comparisons confirmed that ivermectin significantly reduced EV concentration in the 101-150 nm (P < 0.0001), 151-200 nm (P < 0.0001), and 201-250 nm (P < 0.0001) subsets, while other size ranges were not significantly affected. **B:** In the middle intestine, ivermectin (1 µM) significantly inhibited the 101–150 nm, 151–200 nm, 201–250 nm, and 251-300 nm EV subsets released from the middle intestinal region of *Ascaris suum*. Statistical significance is indicated by **** for *P* <0.0001. The EV concentrations were mean ± SEM 1.48 x 10^7^ ± 4.43 ×10^5^ particles mL^−1^ mg^−1^ with 0.01% DMSO and 4.89 x 10^6^ ± 4.08 ×10^5^ particles mL^−1^ mg^−1^ with 1 µM ivermectin in the 101-150 nm EV subsets; EV concentrations were 3.86 x 10^7^ ± 8.76 ×10^5^ particles mL^−1^ mg^−1^ with 0.01%DMSO and 1.41 x 10^7^ ± 8.92 ×10^5^ particles mL^−1^ mg^−1^ with 1 µM ivermectin in the 151–200 nm EV subsets; EV concentrations were 1.30 x 10^7^ ± 3.42 ×10^5^ particles mL^−1^ mg^−1^ with 0.01%DMSO and 5.55 x 10^6^ ± 4.25 ×10^5^ particles mL^−1^ mg^−1^ with 1 µM ivermectin in the 201–250 nm EV subsets; EV concentrations were 3.86 x 10^6^ ± 1.50 ×10^5^ particles mL^−1^ mg^−1^ with 0.01%DMSO and 1.73 x 10^6^ ± 2.18 ×10^5^ particles mL^−1^ mg^−1^ with 1 µM ivermectin in the 251–300 nm EV subsets. Percent reductions in EV concentration (particles mL^−1^ mg ^−1^) following ivermectin treatment are indicated above the red bars. Ivermectin reduced EV concentration (particles mL^−1^ mg ^−1^) by 67% in the 101–150 nm EV size range, 63% in the 151–200 nm range, 57% in the 201–250 nm range, and 55% in the 251–300 nm range. Two-way ANOVA showed a significant inhibitory effect of ivermectin on EV concentration (P = 0.0199; F = 5.51; DFn = 1, DFd = 208), and a significant interaction between the effect of ivermectin and EV size (P < 0.0001; F = 7.52; DFn = 7, DFd = 208), indicating that ivermectin did not reduce all EV size populations equally and had different quantitative effects across EV size groups, consistent with heterogeneity within the intestinal EV population. Post-hoc Šídák multiple comparisons confirmed that ivermectin significantly reduced EV concentration in the 101-150 nm (P < 0.0001), 151-200 nm (P < 0.0001), 201-250 nm (P < 0.0001), and 251-300 nm (P = 0.0008) subsets, while other size ranges were not significantly affected. **C:** In the posterior intestine, ivermectin (1 µM) significantly inhibited EV subsets (101– 150 nm, 151–200 nm, and 201–250 nm) released from the posterior intestinal region of *Ascaris suum*. EV concentrations were 2.16 x 10^7^ ± 9.97 ×10^5^ particles mL^−1^ mg^−1^ with 0.01%DMSO and 6.90 x 10^6^ ± 3.28 ×10^5^ particles mL^−1^ mg^−1^ with 1 µM ivermectin in the 101-150 nm EV subsets; EV concentrations were 5.28 x 10^7^ ± 1.17 ×10^6^ particles mL^−1^ mg^−1^ with 0.01%DMSO and 2.05 x 10^7^ ± 7.25 ×10^6^ particles mL^−1^ mg^−1^ with 1 µM ivermectin in the 151–200 nm EV subsets; EV concentrations were 1.70 x 10^7^ ± 6.05 ×10^5^ particles mL^−1^ mg^−1^ with 0.01%DMSO and 9.11 x 10^6^ ± 6.09 ×10^5^ particles mL^−1^ mg^−1^ with 1 µM ivermectin in the 201–250 nm EV subsets. Percent reductions in EV concentration (particles mL^−1^ mg ^−1^) following ivermectin treatment are indicated above the red bars. Ivermectin reduced EV concentration (particles mL^−1^ mg ^−1^) by 68% in the 101–150 nm EV size range, 61% in the 151–200 nm range, and 46% in the 201–250 nm range. Two-way ANOVA did not show a significant overall main effect of ivermectin across all size subsets (P = 0.8596; F = 0.03; DFn = 1, DFd = 208), but there was a significant interaction between ivermectin treatment and EV size (P < 0.0001; F = 4.89; DFn = 7, DFd = 208). Post-hoc Šídák multiple comparisons confirmed that ivermectin significantly reduced EV concentration in the 101-150 nm (P < 0.0001), 151-200 nm (P < 0.0001), and 201-250 nm (P < 0.0001) subsets, while other size ranges were not significantly affected.

In the anterior intestine, ivermectin reduced the EV concentration (particles mL^−1^ mg ^−1^) by: 69% in the 101–150 nm EV size range; 57% in the 151–200 nm range; and 50% in the 201–250 nm range (Fig. 4A; N=14). Two-way ANOVA showed that there was a significant inhibitory effect (P<0.0001; F=136.45, DFn=1, DFd=208) of ivermectin on the concentration of EVs, as anticipated. There was also a significant interaction between the effect of ivermectin and the EV size (P=0.0013; F=3.53, DFn=7, DFd=208) showing that ivermectin did not reduce all EV size populations equally and had different quantitative effects on the different EV size groups indicating heterogeneity of the EV population.

In the middle intestinal region, ivermectin reduced the EV concentration (particles mL^−1^ mg ^−1^) by: 67% in the 101–150 nm EV size range: 63% in the 151–200 nm range; 57% in the 201–250 nm range; and 55% in the 251–300 nm range (Fig. 4B; N=14). Finally, in the posterior intestinal region, ivermectin reduced the EV concentration (particles mL^−1^ mg ^−1^) by: 68% in the 101–150 nm EV size range: 61% in the 151–200 nm range: and 46% in the 201–250 nm range (Fig. 4C; N=14). Analysis of variance showed that ivermectin significantly reduced the concentrations of the EV produced from the middle and posterior regions of the intestine. As with the anterior region, there were also significant interactions between the size of the EVs and treatment with ivermectin, indicating again that ivermectin does not reduce the concentrations of the different sizes of the EVs equally and indicates heterogeneity of the intestinal EVs.

### 3.4 Proteomic characterization of *A. suum* intestinal EVs

EVs secreted by parasitic nematodes have been demonstrated to contain and deliver bioactive cargo, including small RNAs and proteins, to host cells to modulate innate immune responses (Buck et al., 2014, Eichenberger et al., 2018, Hansen et al., 2019). Having established that the intestine of *A. suum* secretes EVs and that ivermectin inhibits their release, we investigated the protein composition of intestinal EVs to determine whether immune-associated proteins are present and if they were modulated by the effects of ivermectin.

We performed a proteomic analysis on isolated EVs from the whole intestine of *A. suum*. After removal of contaminants and the Pierce Peptide Retention Time Calibration mixture standard entries, we identified 1,574 *A. suum* proteins within intestinal EVs (Supplementary Data File S1), which included classic EV markers, such as Syntenin-1, heat shock proteins (HSP70 and HSP90), annexins, galectins, and flotillins (Flotillin-1 and -2), providing additional support for characterizing these vesicles as EVs. (Buck et al., 2014, Cwiklinski et al., 2015, Kugeratski et al., 2021, Manikantan et al., 2024, Welsh et al., 2024). Using a priority-based functional classification, 96 proteins (6.1%) were annotated with putative immune-related functions, and 130 proteins (8.3%) were associated with digestive functions. The remaining proteins were distributed across major functional categories including translation, protein folding and transport, energy metabolism, and proteolysis (Fig. 5A). A substantial fraction of proteins lacked an informative Gene Ontology Biological Process annotation, consistent with the incompletely characterized *A. suum* proteome. The presumed immune-related repertoire was dominated by heat-shock proteins, transthyretin-like proteins, lectins/galectins, thioredoxins/peroxiredoxins, cathepsins, cyclophilins, and the ASABF antimicrobial peptides (Fig. 5B). The digestive-related repertoire was dominated by proteases/peptidases and glycosidases (Fig. 5C).

**Fig. 5.**
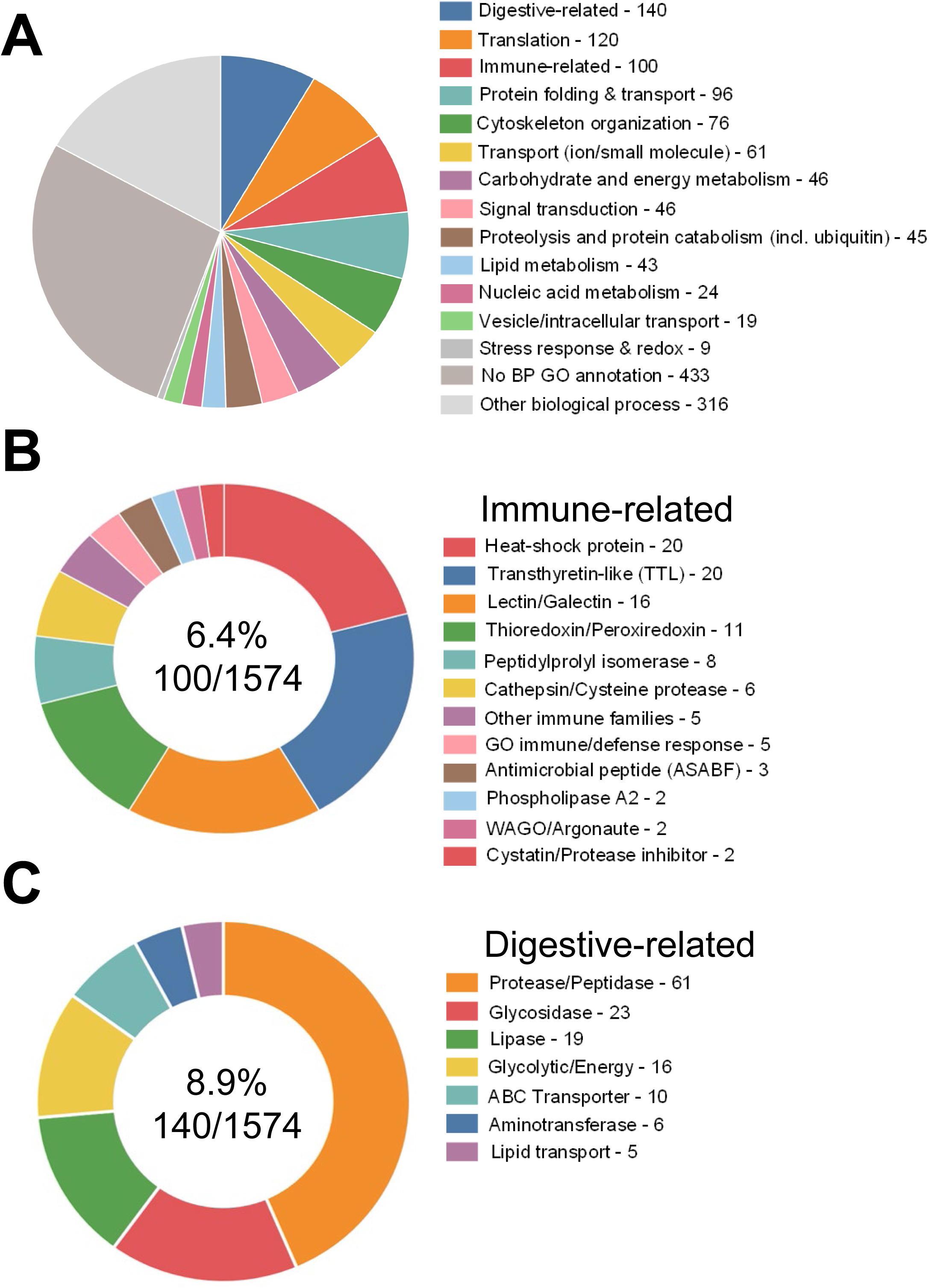
Functional classification of the *A. suum* intestinal-EV proteome. Proteins were classified by a priority scheme: proteins matching curated putative immune-related or digestive-related protein families (by protein name or Gene Ontology annotation) were assigned to those categories first, and all remaining proteins were grouped by Gene Ontology Biological Process (UniProtKB). **A:** Distribution of the 1,574 *A. suum* intestinal-EV proteins across functional categories. Categories and their proportions of the total proteome were: No GO annotation, 436 (27.7%); Other biological process, 265 (16.8%); Digestive-related, 130 (8.3%); Translation, 119 (7.6%); Protein folding & Transport, 116 (7.4%); Immune-related, 96 (6.1%); Carbohydrate & Energy metabolism, 85 (5.4%); Cytoskeleton organization, 68 (4.3%); Proteolysis & Protein catabolism (incl. ubiquitin), 67 (4.3%); Lipid metabolism, 50 (3.2%); Signal transduction, 49 (3.1%); Vesicle/Intracellular transport, 32 (2.0%); Transport (ion/small molecule), 27 (1.7%); Nucleic acid metabolism, 25 (1.6%); and Stress response & Redox, 9 (0.6%). Presumed immune-related and digestive-related proteins form distinct fractions alongside major housekeeping categories such as translation, protein folding/transport, and energy metabolism. Proteins lacking an informative GO Biological Process annotation are shown separately (“No GO annotation” and “Other biological process“), consistent with the incompletely characterized *A. suum* proteome. **B:** Sub-family composition of the 96 putative immune-related proteins; the center value indicates their proportion of the total proteome (6.1%). **C:** Sub-family composition of the 130 digestive-related proteins; the center value indicates their proportion of the total proteome (8.3%).

To provide functional context for the potential immunomodulatory protein candidates that were identified, we annotated each candidate against the primary literature. For each of the major proteins identified, we prioritized direct functional evidence in either *A. suum* or in the human parasite *A. lumbricoides*. If no evidence was present for *Ascaris,* we determined functional evidence using the closest homolog from other nematode species. A list of this evidence can be found in Supplementary Table 3. These results demonstrate that the EVs from the intestine contained a diverse repertoire of proteins associated with different biological processes, including digestion, cellular signaling, and the potential to modulate the host immune system.

### 3.5 Ivermectin alters the immune and digestive related protein cargo of intestinal EVs

Having characterized the protein composition of intestinal EVs, we determined whether the reduction of EV release by ivermectin is accompanied by changes in the EV protein cargo. We compared the protein content of intestinal EVs from ivermectin-treated and DMSO-control treated worms. We collected the 1,574 quantified proteins, and 38 were differentially abundant (P < 0.05; log₂ fold change > 1), with 15 increased and 23 decreased following ivermectin exposure (Fig. 6 & 7).

**Fig. 6.**
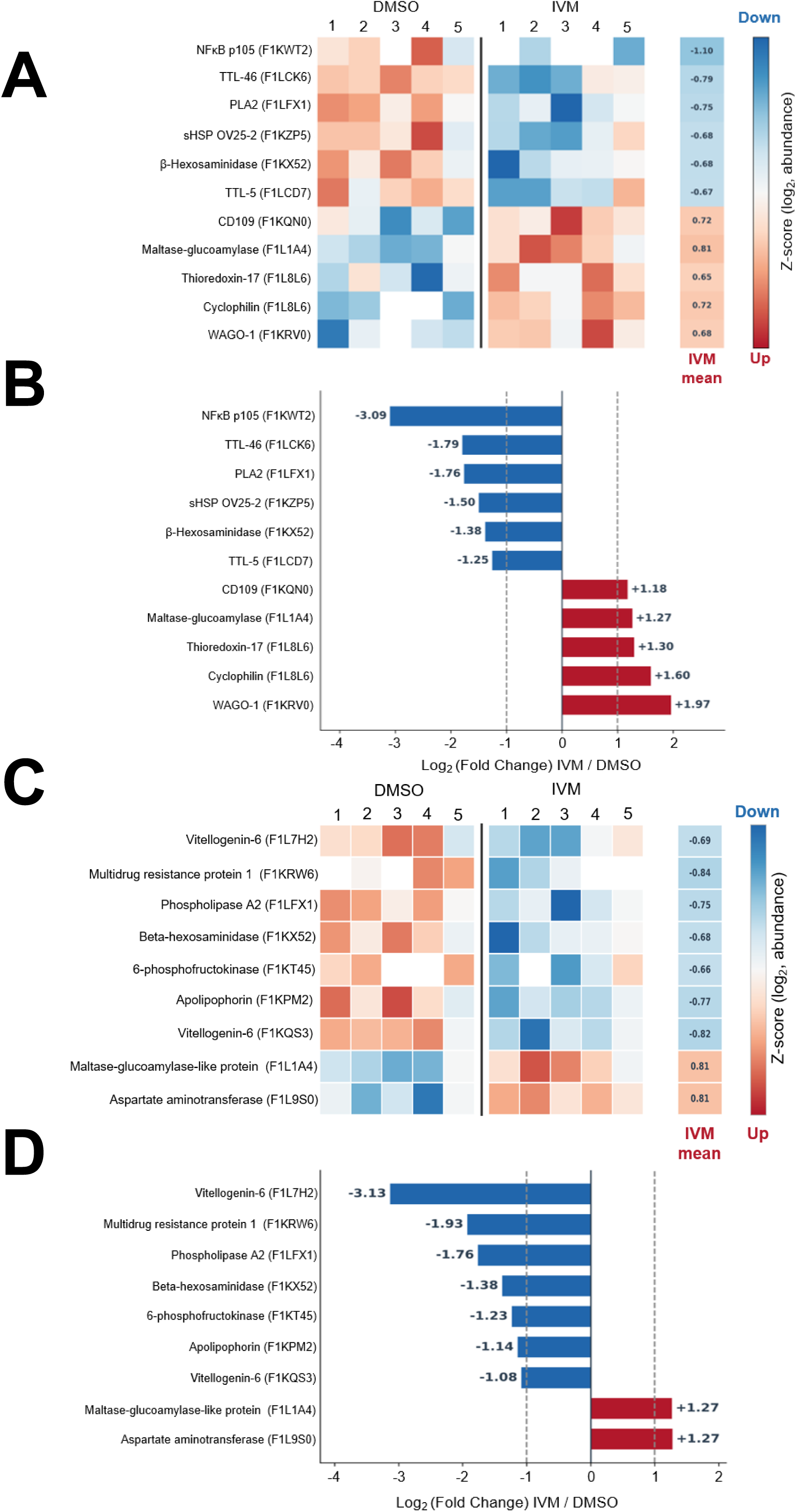
Potential Immune-related and digestive-related proteins significantly affected by ivermectin. Heatmaps of Z-scored log₂ protein abundance and corresponding log₂ fold-change (IVM/DMSO) bar charts for the top 11 potential immune-related and 9 potential digestive-related proteins. Color scale: blue, lower abundance (down); red, higher abundance (up) in 1µM ivermectin compared to the 0.01%DMSO. The proteins are ordered from most downregulated to most upregulated. Ivermectin sample blocks are separated by a vertical divider. The right-hand column shows the mean ivermectin Z-score for each protein. Dashed lines mark log₂FC = 1. Proteins are labeled by name and UniProt accession. All proteins shown are significant (p < 0.05, log₂FC > 1). **A:** Heat map showing the decreased and increased abundance of the top potential immune-related proteins. **B:** log₂ (fold-change) IVM/DMSO bar charts for the top potential immune-related proteins. **C:** Heat map showing the decreased and increased abundance of the top digestion-related proteins. **D:** log₂ (fold-change) IVM/DMSO bar charts for the top digestion-related proteins.

**Fig. 7.**
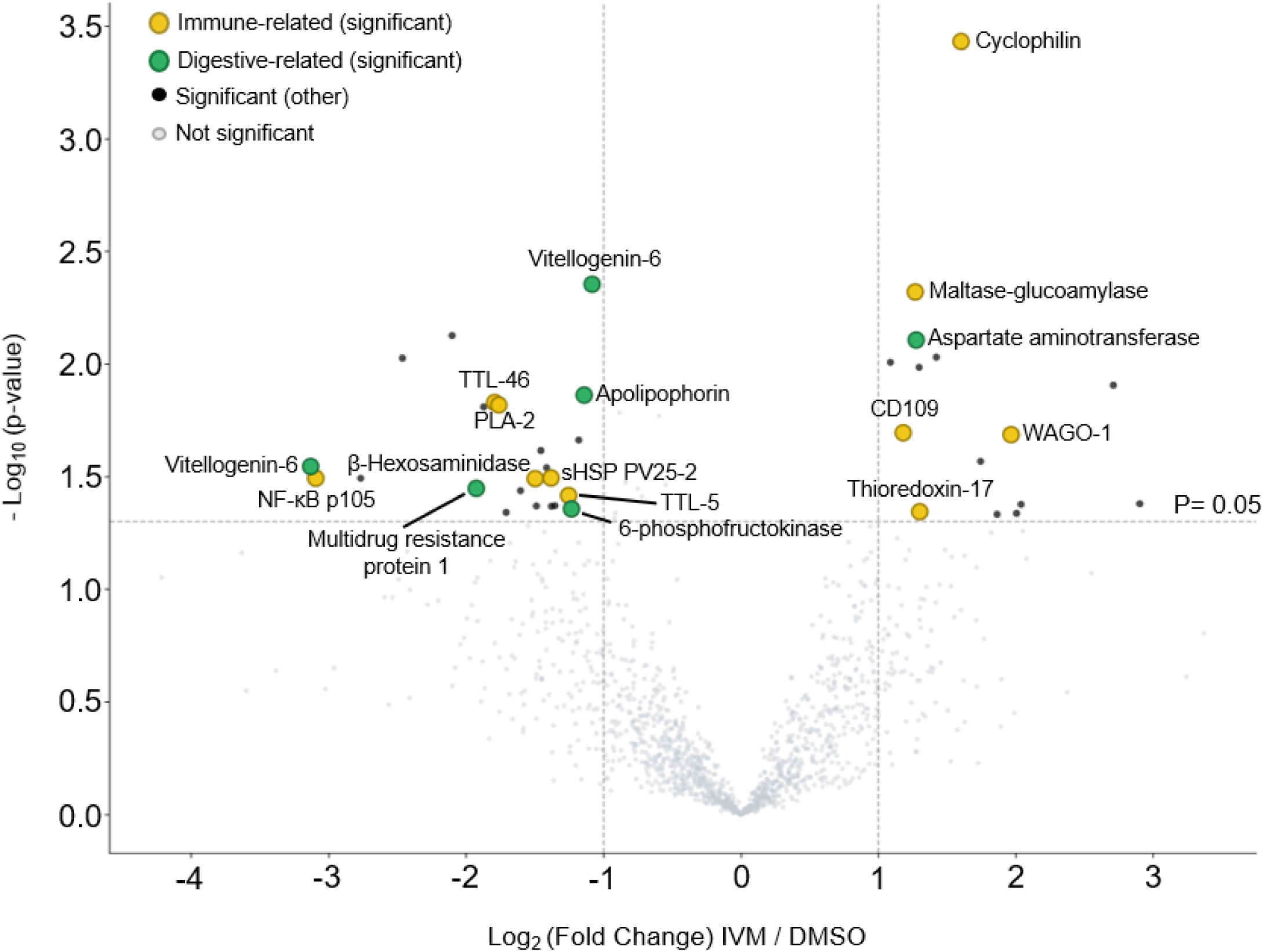
Volcano plot of differential protein abundance by ivermectin treated *A. suum* intestinal EVs. Each point represents one protein; the x-axis shows log₂ (fold change) IVM/DMSO, and the y-axis shows −log₁₀(p-value) (Welch’s t-test, N = 5). Dashed lines indicate the significance thresholds (p = 0.05 and │log₂FC │> 1). Of 1,574 quantified proteins, 38 were significantly altered (23 decreased and 15 increased following ivermectin exposure). Light-grey points, not significant; black points, significant proteins not assigned to the immune or digestive categories; yellow points, significant potential immune-related proteins; green points, significant digestive-related proteins. Significant potential immune-related and digestive-related proteins are labelled by name. Proteins on the left are decreased in ivermectin relative to DMSO. Proteins on the right side of the plot are increased in ivermectin relative to DMSO.

Among the differentially abundant proteins, multiple potential immune-related proteins were altered (Fig. 6A & B). The NF-κB subunit p105, transthyretin-like proteins (TTL-46 and TTL-5), phospholipase A2, and the small heat-shock antigen OV25-2 were decreased following ivermectin exposure, whereas cyclophilin, thioredoxin-17, and the Argonaute protein WAGO-1 were increased. Several digestive-related proteins were also affected (Fig. 6C & D), including vitellogenin-6, multidrug-resistance protein 1, and apolipophorin, which were decreased, and aspartate aminotransferase and a maltase-glucoamylase-like protein, which were increased. These findings indicate that ivermectin inhibits intestinal EV release and alters the abundance of the immune and digestion related protein cargos carried within the vesicles. These observations are consistent with a potential role for intestinal EVs to modulate the host–parasite interaction and identifies that changes in EV protein cargo are an effect of ivermectin interaction on the parasite intestine.

### 3.6 Identification of GluCl channel subunits message in intestine

IVM is a known positive allosteric modulator of glutamate chloride gated channels (GluCls), which are expressed in the neuronal and muscle tissues of nematodes including *A. suum* (Cully et al., 1994, Martin, 1996, Martin et al., 1997). We hypothesized that the mode of action of ivermectin on the intestine was due to the presence of GluCls in the intestine. We used the nucleotide sequences of *glc-2, glc-3, glc-4* and *avr-14* transcripts from *Caenorhabditis elegans* as queries to search the *A. suum* genome (PRJNA62057, version WBPS19) using BLASTN and identified orthologues for all four GluCl subunits (Supplementary Table 1. To determine if *Asu-glc-2, Asu-glc-3, Asu-glc-4,* and *Asu-avr-14* message are expressed in the intestine, we performed RT-PCR on paired intestine and body wall tissues (body wall included nervous and muscle tissues) from five individual female worms. We detected the presence message of all four subunits in both intestinal and body wall cDNA pools, with *Asu-gapdh* serving as a positive control (Fig. 8A – D).

**Fig. 8.**
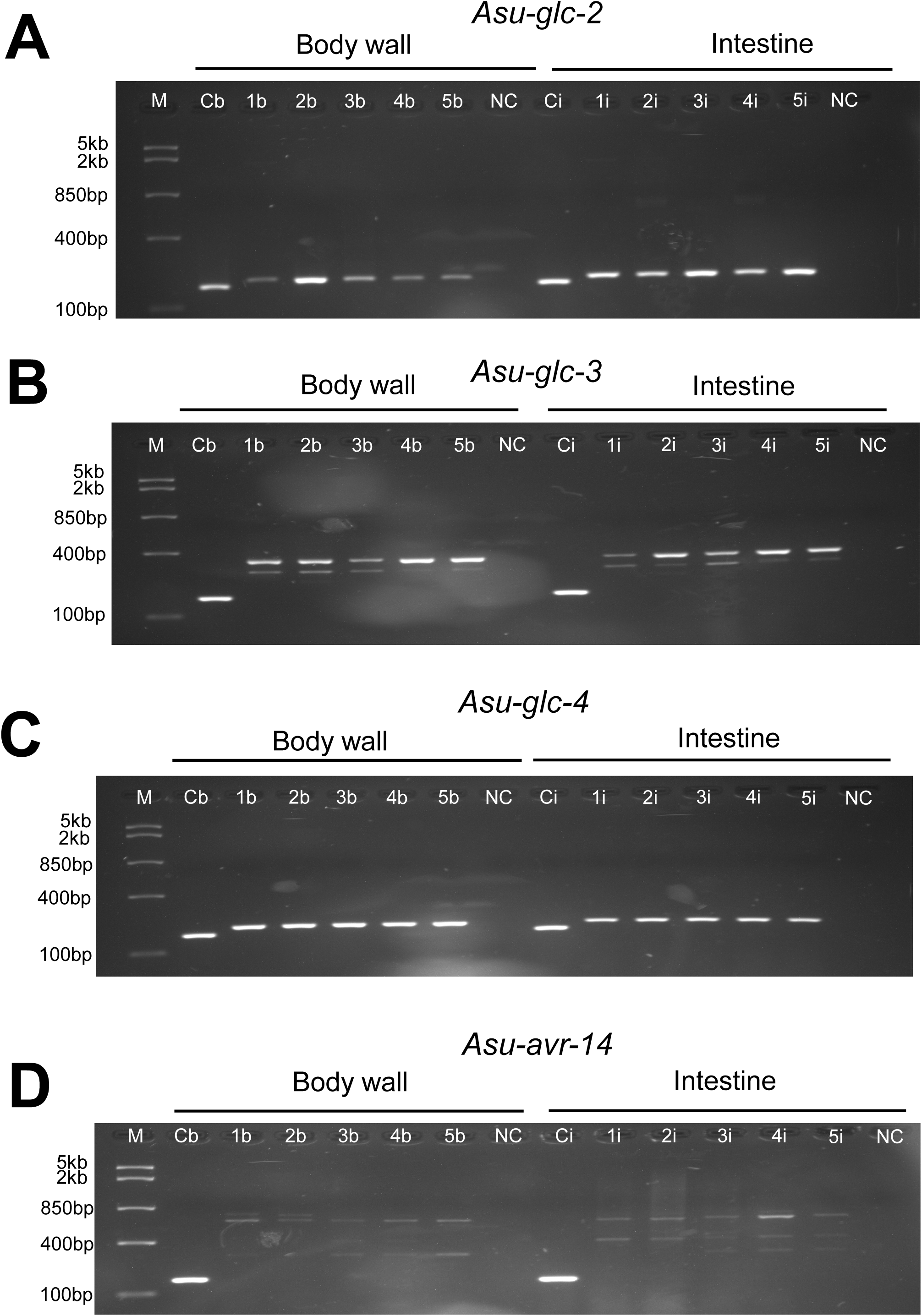
Identification of glutamate-gated chloride (GluCl) channels in the body wall and intestine of adult female *Ascaris suum*. RT-PCR was performed on paired body wall (1b, 2b, 3b, 4b, 5b) and intestine (1i, 2i, 3i, 4i, 5i) samples from five individual adult female worms. *Asu-gapdh* expression in body wall (Cb) and intestine (Ci) served as positive control. NC = negative control (no cDNA template). M = Fast Ruler Middle Range DNA Ladder (Thermo Fisher Scientific). All gel images are cropped; uncropped original images are provided in Supplementary Fig. S4. Images were acquired under UV illumination with an exposure time of 3 s per frame. Amplified products are shown. **A:** *Asu-glc-2* **B:** *Asu-glc-3* **C**: *Asu-glc-4* **D**: *Asu-avr-14*.

Although we confirmed the presence of all four GluCl subunits, only *Asu-glc-2* and *Asu-glc-4,* provided singular distinct bands. For *Asu-glc-3* and *Asu-avr-14* we detected multiple bands suggesting that multiple isoforms of these two subunits may be expressed in the intestine. According to the WormBase parasite genomic database *Asu-glc-3* is predicted to express 9 distinct isoforms. To determine which isoforms of are expressed, we performed RT-PCR with primer sets that target and help determine which specific isoforms are expressed (Fig. 9A & Table 1). Using our isoform specific primers, we found that the intestine expresses at least 3 isoforms of *Asu-glc-3* including isoform 6, 7 and 8 based on product size Fig. 9B. These findings suggest that at least three isoforms of *Asu-glc-3* are expressed in the intestine.

**Fig. 9.**
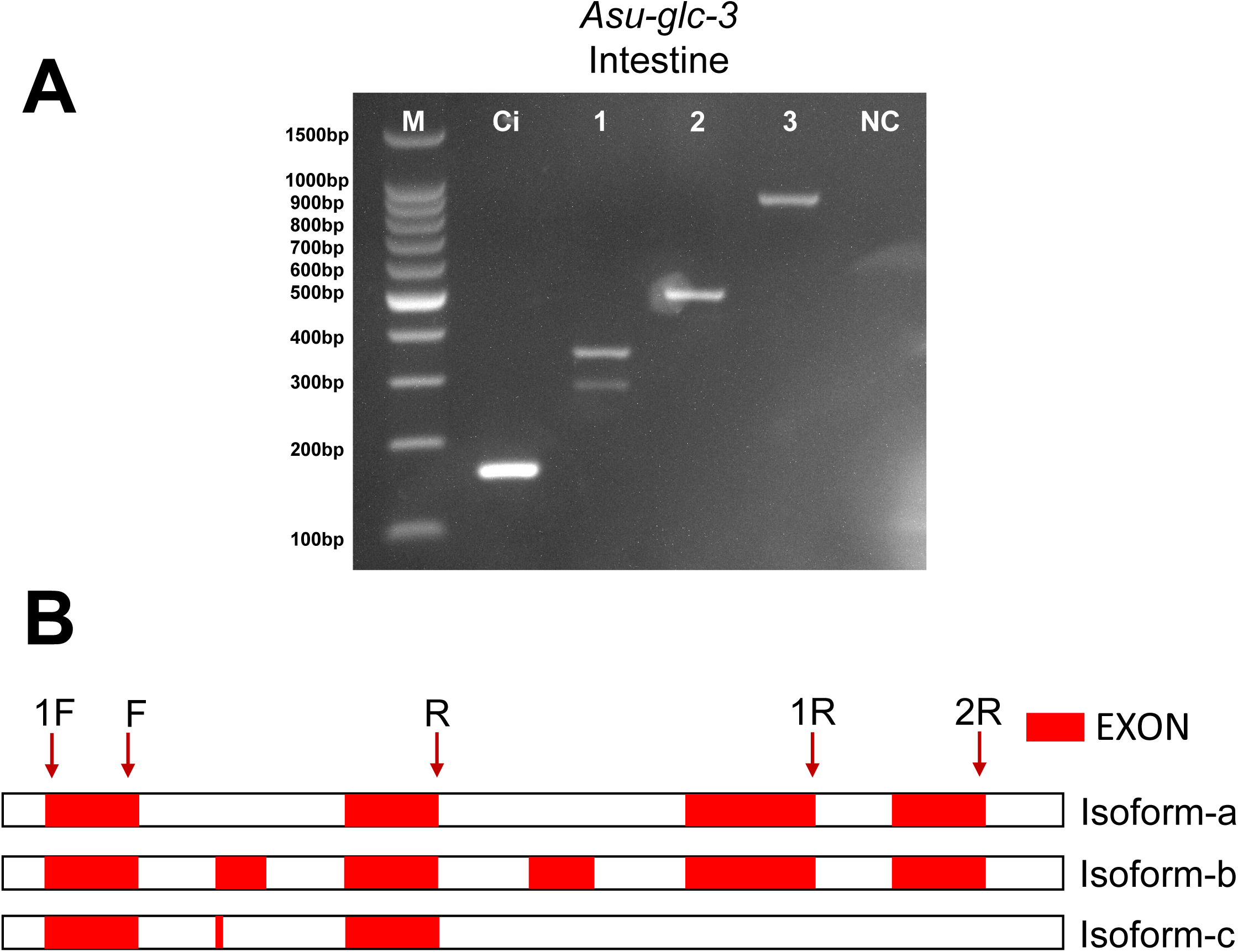
Identification of *Asu-glc-3* isoforms in the intestine of adult female *Ascaris suum*. **A:** Agarose (2%) gel picture of PCR products highlighting *Asu-glc-3* isoforms in the intestine (1, 2, 3) of *Ascaris suum*. DNA ladder is 100bp DNA ladder, Ci= positive control (gapdh), and NC = negative control (no cDNA template). **B:** graphs show the exon size difference between three isoforms. F for each primer set represents forward primer, R,1R and 2R represent reverse primers. The red box represents the exon placed for each isoform. Uncropped original gel images are provided in Supplementary Fig. S4 (E).

**Table 1.** Primer sets used for amplification of *Asu-glc-3* isoforms. The top row of Table 1 represents *Asu-glc-3* isoforms. The leftmost column of Table 1 is three sets of primers. Size of each isoform product length is indicated for each primer pair. Exon sizes for each *glc-3* isoform are listed, corresponding to the amplicons generated by each primer group, based on WormBase Parasite annotations.

| Isoforms<br>Primers |  | 9 | 8 | 7 | 6 | 5 | 4 | 3 | 2 | 1 |
| --- | --- | --- | --- | --- | --- | --- | --- | --- | --- | --- |
|  |  | Exon size | Exon size | Exon size | Exon size | Exon size | Exon size | Exon size | Exon size | Exon size |
| 1 | glc-3 F + R | 352 | 352 | 352 | 352 | 352 | 235 | 235 | 235 | 283 |
| 2 | glc-3 1F + 1R | 378 | 504 | 504 | 504 | 378 | 378 | 378 | 378 | 0 |
| 3 | glc-3 1F + 2R | 785 | 911 | 911 | 911 | 785 | 485 | 485 | 485 | 0 |

To confirm the identity of *Asu-avr-14*, we were unable to generate isoform specific primers due to the large number of predicted isoforms. Therefore, to test for the presence of *Asu-avr-14,* we performed RT-PCR amplifying the full-length *avr-14* transcript from the intestine cDNA pool and performed bidirectional Sanger sequencing. We extracted the product from the gel and performed full-length sequencing and compared the sequence to the database (Supplementary Fig. S3). Our *Asu-avr-14* product was 99% similar to the database suggesting that *Asu-avr-14* is expressed in the intestine of *A. suum* and contributes to a GluCl target of ivermectin.

## 4. Discussion

The intestine of nematodes has often been overlooked as an important anthelmintic target, despite the essential roles it plays in 1) nutrient digestion and absorption, 2) xenobiotic metabolism, 3) innate immunity, and 4) excretion. Here, we have demonstrated that the intestine is a site of EV release and that these EVs contain protein cargo associated with diverse functions, including potential modulation of host immune responses. This EV release is inhibited by ivermectin, an anthelmintic that is typically associated with paralyzing parasites by targeting GluCls on neuromuscular tissues. The inhibition of EV release from the intestine is an additional site of action on the parasite limiting suppression of the host immune systems and allowing it to counter the parasite. There was also an inhibition of the cargo associated with digestion suggesting effects of ivermectin on digestion of the parasite as well.

### 4.1 The parasite intestine is a source of EV release

Moreno *et al*. (2010) proposed the excretory/secretory (ES pore) apparatus as the source of EV release from filarial nematodes; Koehler *et al*. (2021) supported the ES pore as one site of release in *A. suum* by sealing the mouth and anus and leaving only the ES pore open as the only source Graphical Abstract (route 1). Another source of EV release has been identified as the nematode intestine: Buck *et al*. (2014) using *Heligmosomoides polygyrus* demonstrated with electron microscopy that the intestine of secretes EVs from the apical surface of the intestine; Hansen *et al*. (2019) demonstrated that: 1) EVs from *A. suum* derived from intestinal tissue could be released from the anal pore, and 2) EVs from *A. suum* may also be derived in the peri-enteric body fluid (presumably from somatic muscle cells and hypodermis) and released through the ES pore . We found in our studies the EVs, 50-400 nm (Fig. 4) were produced from dissected, isolated *A. suum* intestines and were therefore a source of EVs to be released from the anal pore.

### 4.2 Regional differences in EV release and the effect of ivermectin

The anterior intestinal region of *A. suum* released the greatest number of EVs, suggesting regional specialization in secretory activity. Previous transcriptomic analyses of the contiguous anterior, middle, and posterior regions of the *A. suum* intestine demonstrated longitudinal compartmentalization of gene expression and predicted miRNA regulation, with the anterior region displaying the most distinct molecular and functional profile (Gao et al., 2017). The anterior intestine is thought to be a particularly active absorptive region suggesting that it has more membrane trafficking, endocytosis, exocytosis, and membrane turnover than the other regions of the intestine. The anterior intestine of *A. suum* is also the closest to newly ingested intestinal fluid from the host.

A simplified biogenesis-based classification of EVs often groups them into exosome-like and microvesicle-like populations (Clancy et al., 2021, Welsh et al., 2024). Exosome-like EVs, approximately 30-50 nm, form as intraluminal vesicles within cytoplasmic multivesicular bodies and are released upon fusion with the plasma membrane. In contrast, microvesicle-like EVs, typically 100-1000 nm, are produced from direct outward budding and fission of the plasma membrane (Clancy et al., 2021). The anterior, middle, and posterior regions of the intestine displayed a wide size distribution with most 50-300 nm. The peak sizes from the three regions were 151-200 nm (Fig. 2), suggesting that these and the bigger EVs were microvesicles produced by membrane blebbing of the enterocytes of the intestine. Interestingly large microvesicle-like EVs are also produced by *A. suum* body muscle cells (Martin et al., 1990).

Microvesicle formation at the plasma membrane surface relies on Ca^2+^ dependent remodeling, actomyosin contraction, Rho-family GTPases, ARF6 signaling and cargo loading(van Niel et al., 2018, Clancy et al., 2021). Ivermectin significantly reduced EV release from all three regions of the intestine and interestingly there was a significant interaction between the effect of ivermectin and the EV sizes. We interpret the significant interaction between ivermectin treatment and EV size as evidence that ivermectin does not uniformly suppress all EV populations because of heterogenous EV release mechanisms. Some (27%) of the EV release was not sensitive to ivermectin. Across all three intestinal regions, the smallest EV size range examined (101-150 nm) showed the greatest proportional reduction (67%-69%), with inhibition decreasing progressively in larger size categories (46%-57% in the 201-250 nm). This gradient suggests that ivermectin selectively inhibits the release of smaller EV populations.

We have demonstrated that the intestine expresses GluCl subunit message including *Asu-glc-2, Asu-glc-3, Asu-glc-4,* and *Asu-avr-14* GluCl and that ivermectin inhibits the release of EVs with an *IC50* 64nM. Activation of GluCl ion-channels by ivermectin would allow entry of Cl^−^ ions, hyperpolarizing the enterocytes and reducing the entry of extracellular Ca^2+^ and the release of intracellular calcium which is required for vesicle trafficking and exocytosis (Cully et al., 1994, Savina et al., 2003, Atif et al., 2017, Loghry et al., 2020). The suppression of calcium-dependent secretory signaling in enterocytes is therefore plausible mechanism linking GluCl activation to reduced EV release.

### 4.3 Ivermectin alters the putative immune-related protein cargo of intestinal EVs

Proteomic analysis identified 1,574 proteins in *A. suum* intestinal EVs, including 96 (6.1%) associated with putative immune-related functions and 130 (8.3%) with digestive functions, Fig. 5. The presence of potential immunomodulatory proteins including heat-shock proteins, transthyretin-like proteins, lectin/galectins, cathepsin/cysteine proteases, cyclophilins and WAGO/Argonaute proteins supports the hypothesis that these EVs carry a cargo that have a role for these vesicles in parasite–host cross talk limiting the host immune response to the parasite (Hansen et al., 2019). Comparison of the ivermectin treated intestinal EVs with the control DMSO treated intestinal EVs, showed that ivermectin not only reduced the number of EVs released from the intestine but also altered their protein cargo: 38 proteins were differentially abundant, including several potential immune-related proteins (Fig. 6 & 7)

Some of these ivermectin-downregulated proteins that have defined immunomodulatory are discussed below. Their roles illustrate how the reduction of the EV cargo could restore host immune recognition. Phospholipase A2 (PLA2), which was downregulated, hydrolyzes membrane phospholipids to release arachidonic acid for the production of eicosanoids, including prostaglandins (Dennis et al., 2011). The prostaglandins are lipid mediators involved in inflammatory and immune regulation (Ricciotti and FitzGerald, 2011). In parasitic infections, parasite and host derived prostaglandins have been implicated in host–parasite interactions and immune modulation (Brattig et al., 2006, Kubata et al., 2007, Maizels and McSorley, 2016). A reduction of PLA2 in ivermectin-treated EVs is anticipated to diminish the parasite’s capacity to manipulate host eicosanoid-dependent inflammatory signaling.

Transthyretin-like proteins were also reduced significantly, including TTL-46 and TTL-5. These proteins belong to a conserved nematode-specific protein family that has been identified in excretory/secretory products from *A. suum* and other parasitic nematodes (Hewitson et al., 2008, Wang et al., 2013, Tritten et al., 2021). Although their precise functions remain to be fully understood, their presence in parasite secretomes implies involvement in host–parasite interactions (Tritten et al., 2021). The reduction in EVs of this conserved secretome-associated protein family indicates a change in the host-interface proteome interaction. The small heat-shock protein OV25-2-like antigen belongs to a broader group of nematode small heat-shock proteins, including stage-specific excretory/secretory sHSPs that are immunogenic and have been implicated in interactions with host intestinal mucosal cells (Younis et al., 2011, Pérez-Morales and Espinoza, 2015). Its reduction following ivermectin exposure indicates a reduction of immunogenic stress-associated EV cargo. There is also a reduction of NF-κB1 p105 that corresponds to a key regulator of NF-κB signaling, a pathway centrally involved in innate immune and inflammatory gene expression in metazoan systems (Hayden and Ghosh, 2008, Liu et al., 2017). The decrease in the annotated NF-κB1 p105 protein may indicate altered abundance of immune/signaling-associated cargo. The specific function of NF-κB1 p105 in *A. suum* EVs remains to be determined.

A subset of immune-or stress-associated proteins increased following ivermectin treatment, including cyclophilin, thioredoxin-17, and the Argonaute/WAGO protein WAGO-1. Helminth thioredoxin/peroxiredoxin-related antioxidant proteins have been reported in excretory/secretory products that promote alternatively activated macrophage phenotypes and Th2-skewed responses in experimental models (Donnelly et al., 2005, Donnelly et al., 2008). The increase in thioredoxin-17 may reflect an ivermectin-induced redox or stress response rather than enhanced immune suppression. The inclusion of WAGO-1 is interesting because: nematode EVs contain small RNAs together with worm-specific Argonaute proteins; and parasite-derived EV RNAs can modulate host innate immune responses (Buck et al., 2014, Chow et al., 2019). Direct EV small-RNA profiling and functional uptake assays are necessary to determine if ivermectin alters the RNA-mediated immunomodulatory potential of *A. suum* intestinal EVs.

We identified other potential immune-related proteins in the intestinal EVs of *A. suum,* including cystatins, galectins and a serpin, all of which showed some level of IVM modulation but did not reach statistical significance. We also found a MIF/mif-2 ortholog that showed no detectable IVM effect. At list of these proteins can be found in Supplementary Table 3. While we have not directly shown that our identified proteins have immunological effects on the host, other homologues in other parasite species have been demonstrated to have direct modulation on mammalian immune systems. For example, cystatins from *A. lumbricoides*, which is synonymous with *A. suum* (Leles et al., 2012) and the filarial nematode *Acanthocheilonema viteae* attenuate allergic and colitic inflammation and induce IL-10-producing regulatory macrophages (Coronado et al., 2017, Coronado et al., 2019, Schnoeller et al., 2008). In *B. malayi* the galectins Bma-LEC-1 and Bma-LEC-2 induce cell apoptosis of Th1 cells of mice and Bma-LEC-2 increases IL-10 production in human macrophages (Loghry et al., 2022). TsSERP1 (Serpins which are serine protease inhibitors) in *Trichinella spiralis* inhibited human neutrophil elastase and the production of proinflammatory cytokines and chemokines (Kobpornchai et al., 2022). Finally, in *Haemonchus contortus*, the recombinant MIF protein directly binds host goat monocytes and modulates their immune activity by suppressing TNF-α, IL-1β, IL-12p40, MHC-II expression, NO production, and phagocytosis, while upregulating IL-10 and TGF-β secretion (Wang et al., 2017, Karabowicz et al., 2022).

### 4.5 Digestive, lipid-transport, and detoxification-related EV cargo as additional drug-relevant targets

In addition to immune-associated proteins, ivermectin altered EV proteins (Supplementary Data File S1) linked to nutrient handling, lipid transport, and xenobiotic defense, including decreased vitellogenin-6, apolipophorin, and multidrug-resistance protein 1 (MDR-1, a P-glycoprotein). The reduction of MDR-1 is pharmacologically and therapeutically relevant because these nematode proteins are ATP-dependent efflux transporters that reduce ivermectin in the parasite and have been implicated in the resistance of ivermectin and macrocyclic lactone anthelmintic resistance (Ardelli and Prichard, 2013, Janssen et al., 2015, Lespine et al., 2024). Given that MDR-1 and related ABC transporters are established candidate markers of macrocyclic lactone resistance in multiple nematode species, the ivermectin-induced reduction of MDR-1 in intestinal EV cargo raises the possibility that EV-associated efflux transporter levels could serve as a phenotypic indicator of ivermectin exposure or susceptibility at the tissue level, meriting further investigation as a potential biomarker for resistance screening. Other helminth xenobiotic-metabolizing and transport systems, including cytochrome P450s, UDP-glycosyl/glucuronosyltransferases, and ABC transporters, are associated with anthelmintic metabolism, deactivation, efflux, and resistance (Laing et al., 2015, Matoušková et al., 2016, Matoušková et al., 2018).

### 4.6 Tissue-level resolution of immune-active EV cargo

By localizing EV release specifically to the intestine, our study assigns putative immune-related EV proteins to one defined anatomical source rather than to whole-worm ES material alone. This tissue-level resolution matters because it identifies the intestine as a major secretory interface from which immune-active cargo is delivered into the host environment, and because it pinpoints a tissue in which ivermectin both inhibits secretion and remodels cargo. All these results support a model in which *A. suum* intestinal EVs shape host immunity to promote parasite persistence, and in which ivermectin disrupts this process at its anatomical source.

## 5. Conclusion

The present study supports the conclusion that the *A. suum* intestine is a major source of EV release, expresses known ivermectin-targeted GluCl subunits, including *Asu-glc-2*, *Asu-glc-3*, *Asu-glc-4*, and *Asu-avr-14*, and produces EVs that carry cargo proteins with immunological, metabolic, and drug-response relevance. Ivermectin inhibits intestinal EV secretion and alters the abundance of immune-, digestive-, lipid-transport-, and detoxification-associated proteins carried within these vesicles. These findings suggest that part of the antiparasitic activity of ivermectin involves disruption of EV-mediated host– parasite communication and parasite secretory homeostasis, in addition to its effects on neuromuscular and other excitable tissues. The data reinforces the view that the nematode intestine is a functionally important interface for secretion, host interaction, and a drug site of action.

## Author contributions

**Dongjie Liu**: Performed the experiments and acquired majority of the data, provided interpretations and designs of the experiments, wrote the manuscript and designed figures. **Paul Williams** provided interpretations and conceptions of the data, helped design and perform genetic experiments, wrote, and edited the manuscript and helped design the figures. **Michael Kimber** contributed to interpretation of data and methodology. **Alan Robertson** provided interpretations, conceptions, and designs of experimental procedures. **Richard Martin** provided the conceptualization of the project, methodology, funding acquisition, resources, project administration, supervision, and wrote and edited the manuscript. All authors have reviewed the manuscript.

## Acknowledgements

This study was supported by NIH NIAID Grants R01AI047194, R01AI155413, to RJM, the EA Benbrook Endowed Chair of Pathology and Parasitology and by ISU CVM Seed Grant funding from the Dr Stephen G. Juelsgaard Dean’s award to PDW. The funders had no role in study design, data collection and analysis, decision to publish, or preparation of the manuscript. PDW received salary from the NIH Grants. The funding agencies had no role in the design, execution, or publication of this study. The content is solely the responsibility of the authors and does not necessarily represent the official views of the NIH National Institute of Allergy and Infectious Diseases. We acknowledge JBS swift Marshalltown, IA for access for the collection of the *Ascaris suum* used in these experiments.

## Data availability statement

The data that support the findings of this study are available from the corresponding author upon reasonable request.

**Supplementary Table 1. Primer sequences used for RT-PCR**

Forward (F) and reverse (R) primers targeting the glutamate-gated chloride channel subunit genes *Asu-glc-2*, *Asu-glc-3*, *Asu-glc-4*, and *Asu-avr-14*, with the housekeeping reference gene *gapdh* for a positive control. Full-length *avr-14* primers used for Sanger sequencing are denoted FLF (forward) and FLR (reverse).

**Supplementary Table 2. Gene identifiers and accession numbers for genes targeted by RT-PCR.**

WormBase ParaSite gene IDs and corresponding accession number for each of the four glutamate-gated chloride channel subunit genes (*Asu-glc-2*, *Asu-glc-3*, *Asu-glc-4*, and *Asu-avr-14*) targeted by the primer sets in Supplementary Table 1 from the *A. suum* genome PRJNA62057.

Supplementary Table 3. Putative immunomodulatory proteins identified in the *Ascaris suum* intestinal EV proteome.

Functional evidence for potential immune-related proteins either directly from *Ascaris*, characterized homologs in filarial and/or trichinellid nematodes. The corresponding supporting literature is provided.

---

**Supplementary Fig. S1.**
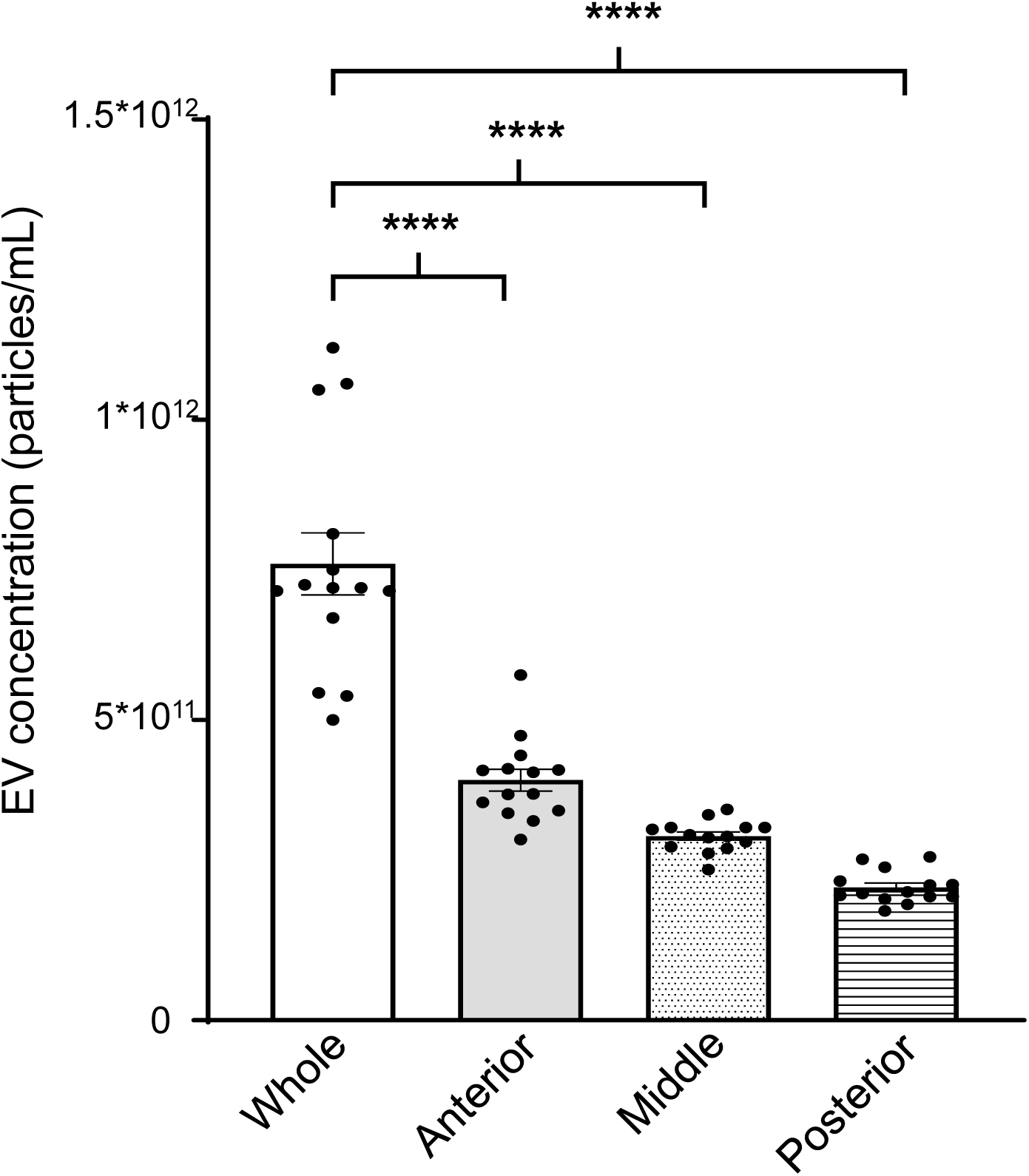
EV concentrations are released from the whole intestine and from anterior, middle, and posterior regions. Quantitative analysis revealed mean concentrations (not normalized to wet weight) of 7.60 x 10^11^ ± 5.16 ×10^10^ particles mL^−1^ from whole intestine, 4.00 x 10^11^ ± 1.84 ×10^10^ particles mL^−1^ from anterior, 3.07 x 10^11^ ± 6.92 ×10^9^ particles mL^−1^ from middle, and 2.21 x 10^11^ ± 7.25 ×10^9^ particles mL^−1^ from posterior intestine. One-way ANOVA model analysis was used to compare the means of EV size profiles following drug treatment with statistical significance determined using post-hoc Šídák test (*P*-values < 0.05 being considered significant). Significant differences in EV release were detected when the whole intestine was compared with the anterior, middle, or posterior intestinal regions. Statistical significance is indicated by **** for *P* <0.0001. Data are presented as mean ± SEM, and *N* represents the number of independent experiments (*N* = 14).

**Supplementary Fig. S2.**
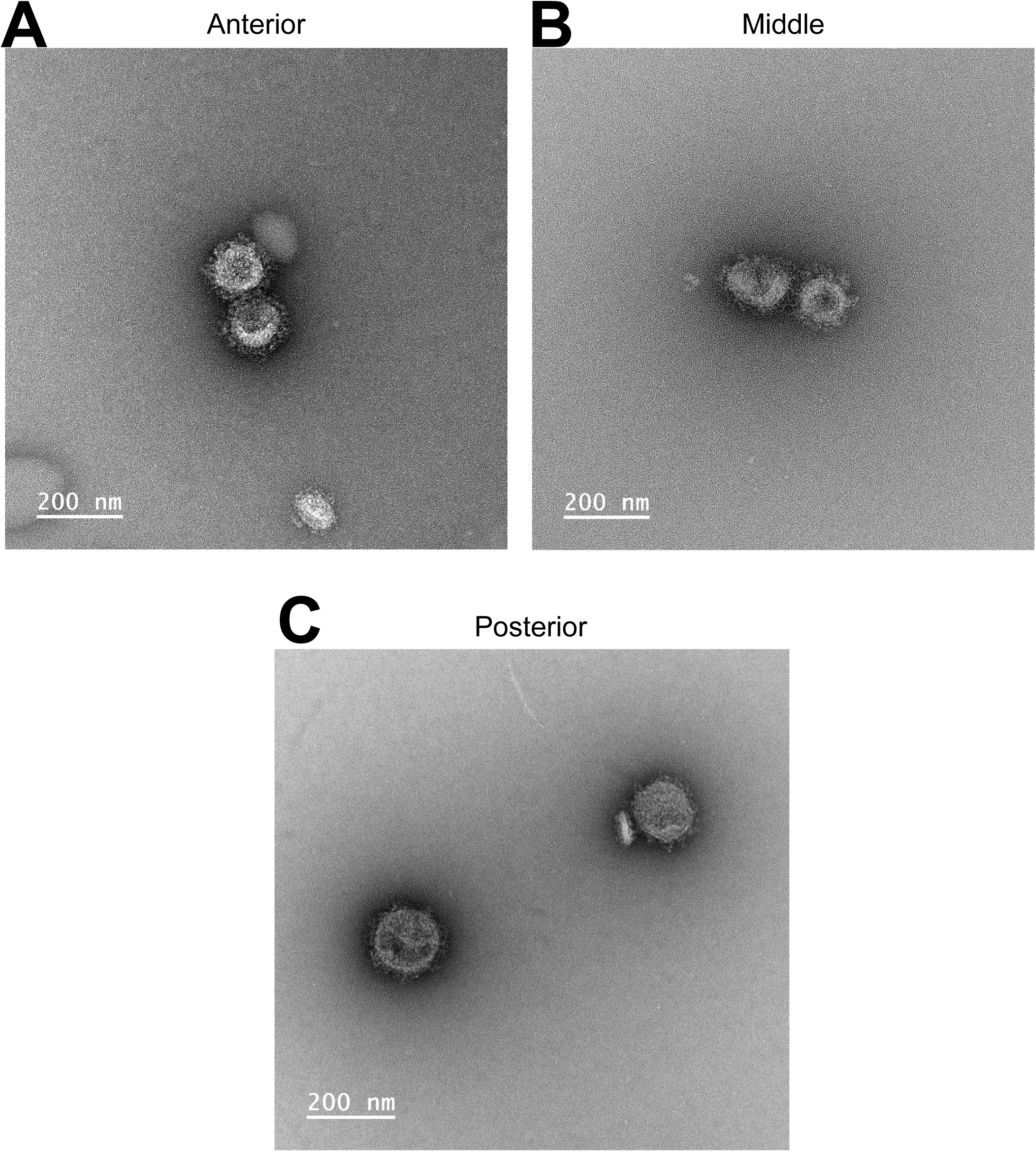
Uncropped TEM images of EVs in anterior, middle and posterior intestines of *Ascaris suum*. Enlarged original uncropped transmission electron microscopy (TEM) images showing EVs (100–200 nm) released into culture media by the anterior, middle, and posterior regions of the intestine from adult female *A. suum* after 4 h incubation in *Ascaris* perienteric fluid containing 0.01% DMSO (pH 7.6). EVs were isolated and purified using ultracentrifugation, SEC columns, and 0.2 µm filtration. Scale bars are indicated.

**Supplementary Fig. S3.**
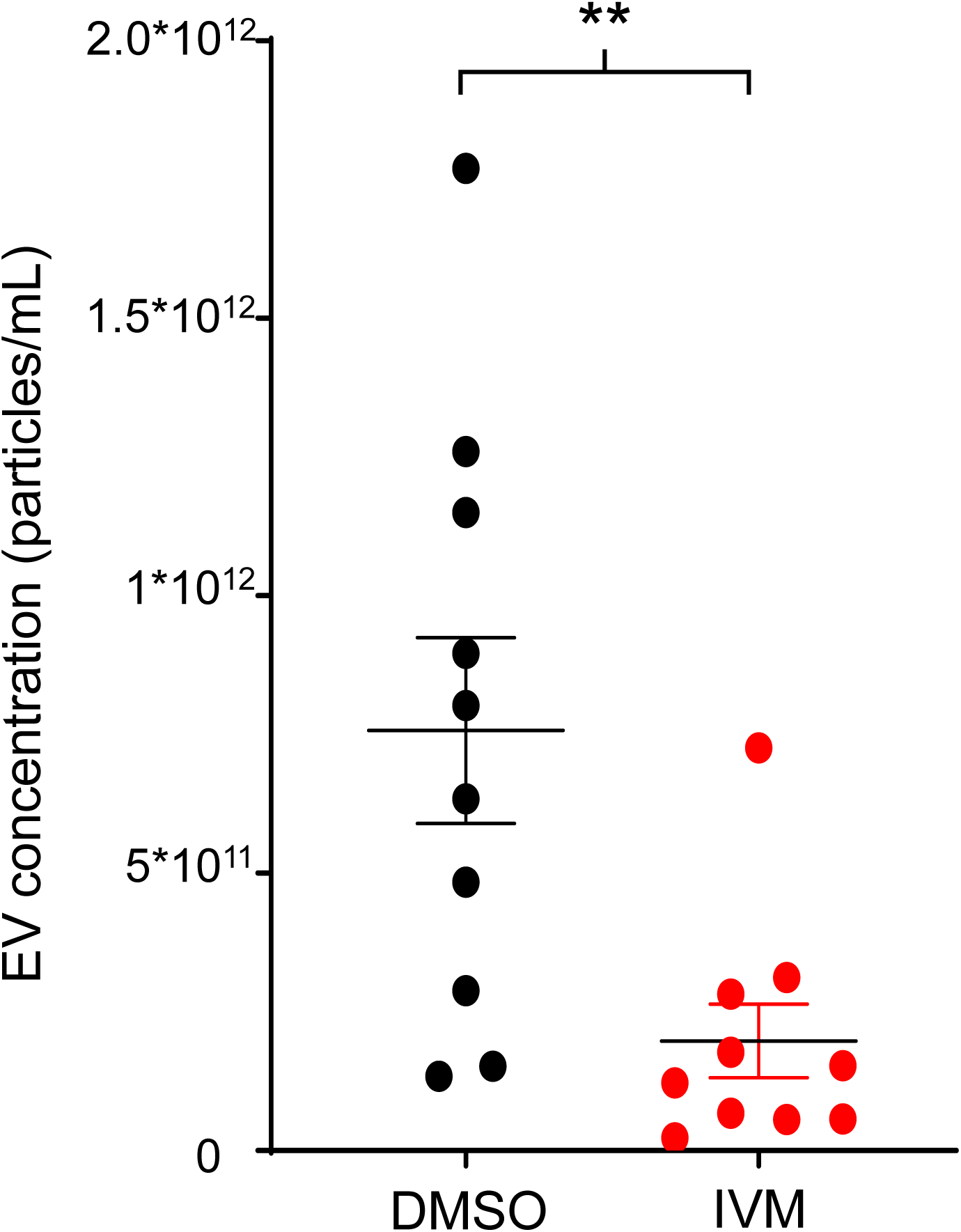
Ivermectin inhibits EV secretion from adult female *Ascaris suum*. Dot plot analysis demonstrates the inhibitory effect of ivermectin (IVM) on extracellular vesicle (EV) secretion. Adult female *Ascaris suum* worms were cultured at 37 °C in Ascaris Ringer solution containing either 1 µM ivermectin or 0.01% DMSO (pH 7.8). Culture media were collected after 24 h, and EVs were isolated and quantified. Statistical comparisons were performed using an unpaired *t*-test, with statistical significance indicated by ** for *P* <0.01 (P=0.0092). EV concentrations were 7.57 x 10^11^ ± 1.67 x 10^11^particles mL^−1^ in the 0.01% DMSO group and 1.98 x 10^11^ ± 6.6 x 10^10^ particles mL^−1^ in the 1 µM ivermectin group from whole adult female worms. Data are presented as mean ± SEM, and *N* represents the number of independent experiments (*N* = 10).

**Supplementary Fig. S4.**
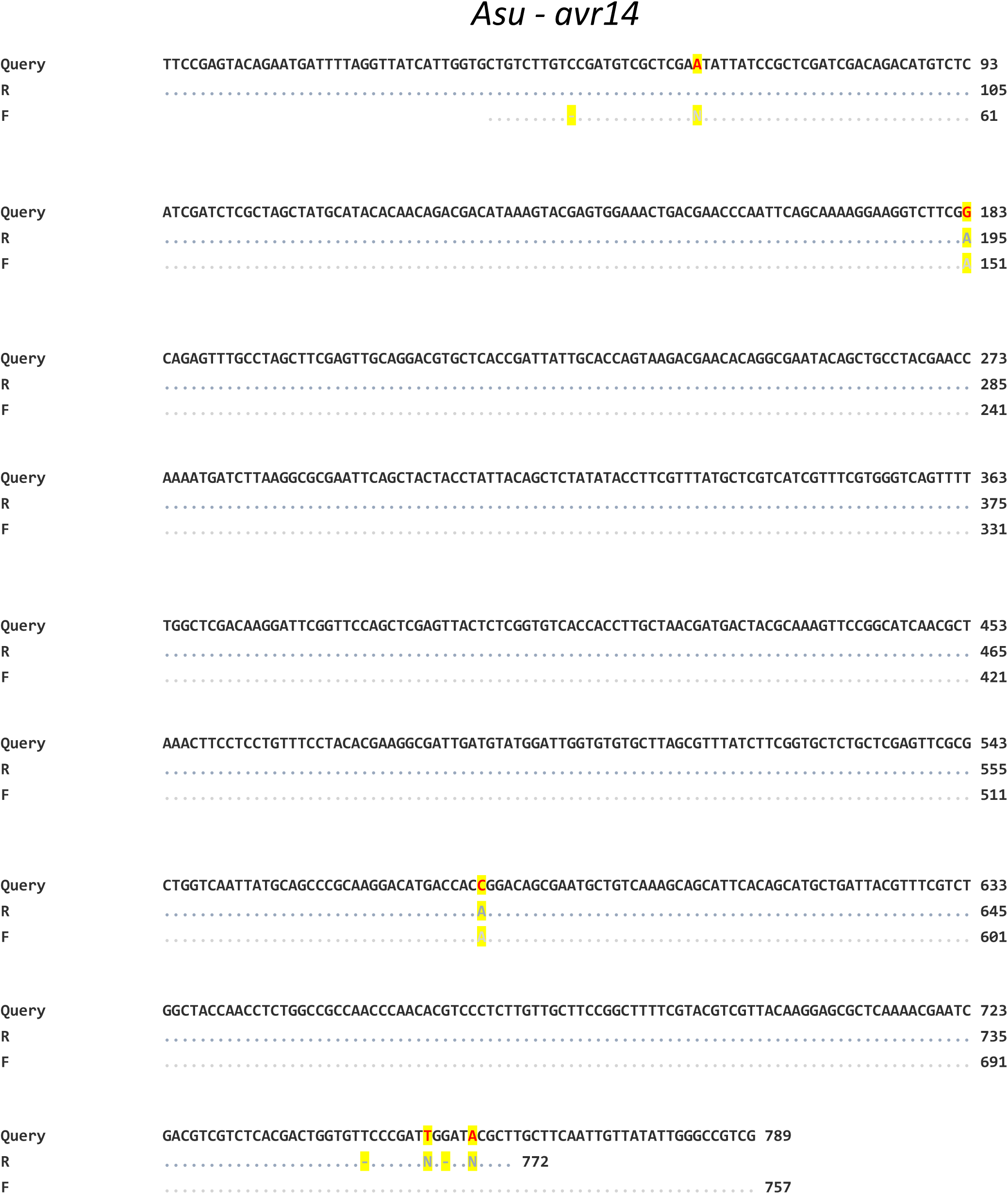
Full-length sequencing of *Asu-avr-14* from the intestine of adult female *Ascaris suum*. RT-PCR was used to amplify the full-length *avr-14* transcript from intestinal tissue. “Query” represents the reference full-length *avr-14* sequence from the database. “R” denotes the *avr-14* product amplified with the reverse primer, and “F” denotes the *avr-14* product amplified with the forward primer. “N “(Ambiguity Code) represents an unknown nucleotide (base); “– “(Gap/Deletion) represents a deletion or a gap in the sequence introduced to align it with other, often longer, sequences (Multiple Sequence Alignment). All other error codes indicate a matching error (mismatch). INSDC Sequence ID: AEUI03000006.1

**Raw_gel Fig. S5.**
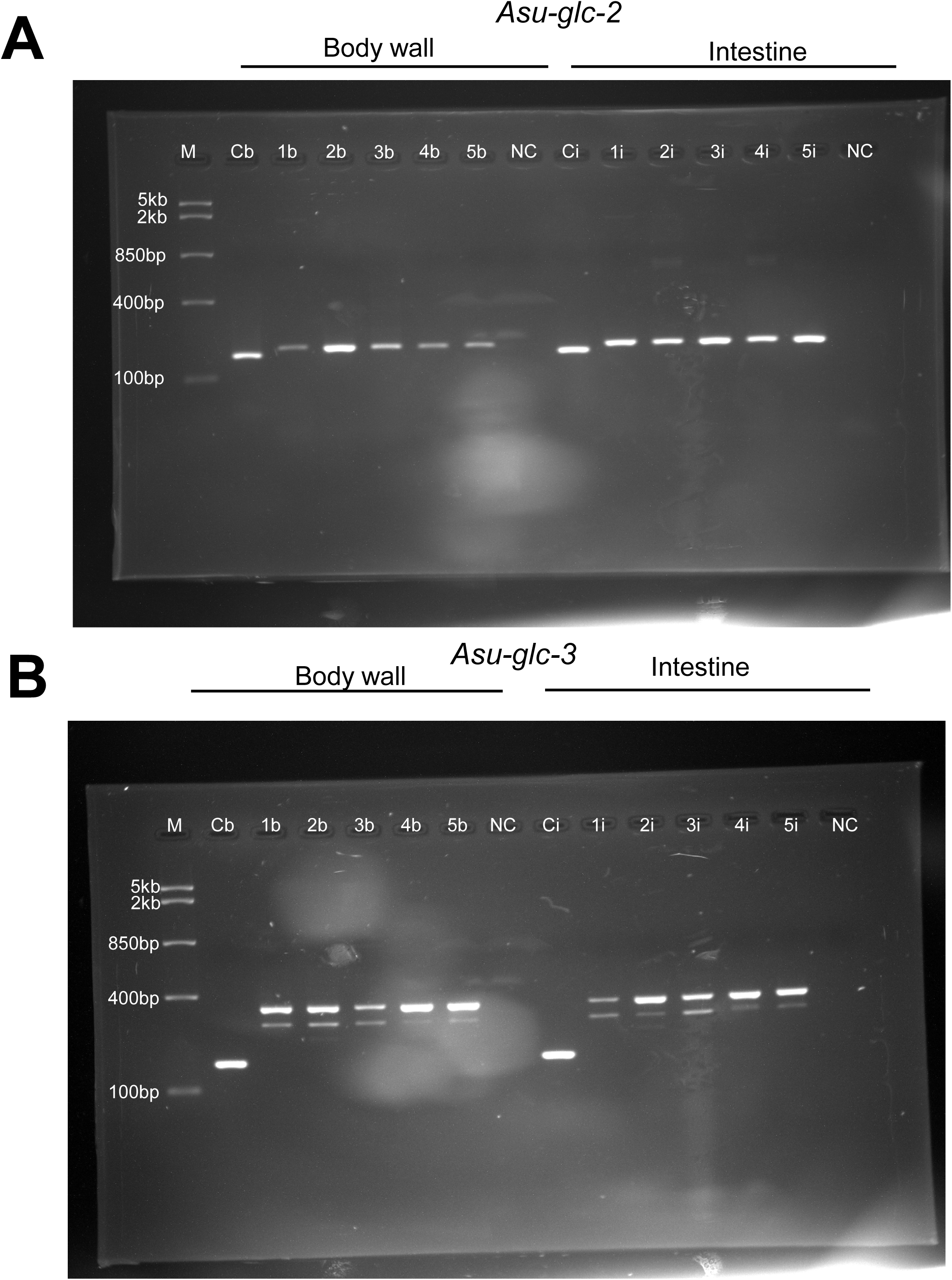
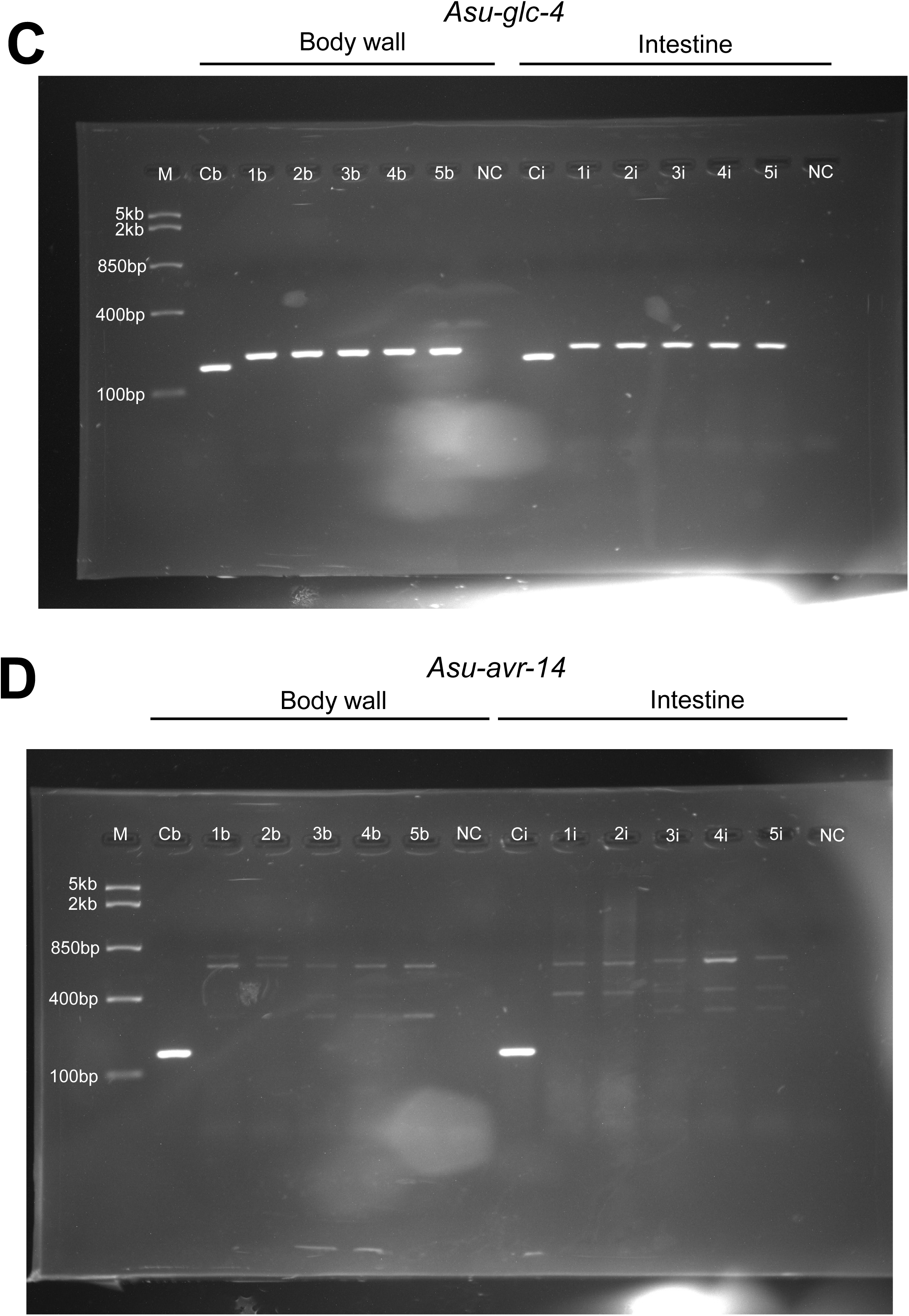
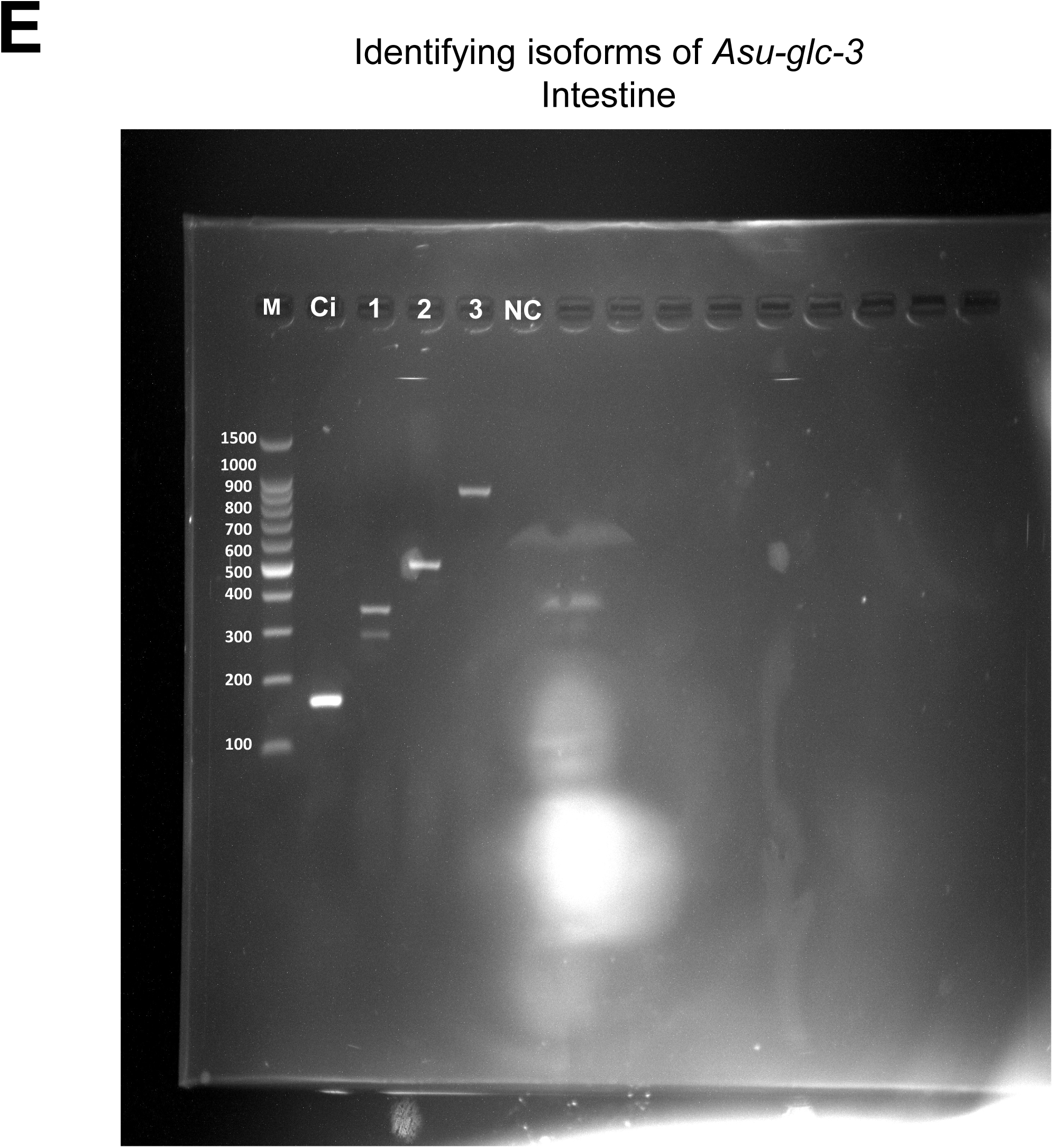
Uncropped images of the Identification of GluCl channel subunits and isoforms in the body wall and intestine of adult female *Ascaris suum*. RT-PCR was performed on paired body wall (1b–5b) and intestinal (1i–5i) samples from five individual adult female worms. *Asu-gapdh* expression in body wall (Cb) and intestine (Ci) served as positive control. NC = negative control (no cDNA template). M = Fast Ruler Middle Range DNA Ladder (Thermo Fisher Scientific). Amplified products are shown for **A:** *Asu-glc-2*, **B:** *Asu-glc-3*, **C:** *Asu-glc-4*, **D:** *Asu-avr-14*, and **E:** *Asu-glc-3* isoforms. Images were captured under UV illumination with an exposure time of 3 s per frame. Uncropped original gel images are presented here.

## References

Ardelli, B. F., Prichard, R. K. 2013. Inhibition of P-glycoprotein enhances sensitivity of Caenorhabditis elegans to ivermectin. Vet Parasitol, 191, 264–75.

Arena, J. P., Liu, K. K., Paress, P. S., Frazier, E. G., Cully, D. F., Mrozik, H., Schaeffer, J. M. 1995. The mechanism of action of avermectins in Caenorhabditis elegans: correlation between activation of glutamate-sensitive chloride current, membrane binding, and biological activity. J Parasitol, 81, 286–94.

Atif, M., Estrada-Mondragon, A., Nguyen, B., Lynch, J. W., Keramidas, A. 2017. Effects of glutamate and ivermectin on single glutamate-gated chloride channels of the parasitic nematode H. contortus. PLoS Pathog, 13, e1006663.

Brattig, N. W., Schwohl, A., Rickert, R., Büttner, D. W. 2006. The filarial parasite Onchocerca volvulus generates the lipid mediator prostaglandin E(2). Microbes Infect, 8, 873–9.

Brownlee, D. J., Holden-Dye, L. C Walker, R. J. 1997. Actions of the anthelmintic ivermectin on the pharyngeal muscle of the parasitic nematode, Ascaris suum. Parasitology, 115 (Pt 5), 553–61.

Buck, A. H., Coakley, G., Simbari, F., Mcsorley, H. J., Quintana, J. F., LE Bihan, T., Kumar, S., ABREU-Goodger, C., Lear, M., Harcus, Y., Ceroni, A., Babayan, S. A., Blaxter, M., Ivens, A. C Maizels, R. M. 2014. Exosomes secreted by nematode parasites transfer small RNAs to mammalian cells and modulate innate immunity. Nat Commun, 5, 5488.

Campbell, W. C. 2012. History of avermectin and ivermectin, with notes on the history of other macrocyclic lactone antiparasitic agents. Curr Pharm Biotechnol, 13, 853–65. CDC. 2024. About Ascariasis [Online]. Available: https://www.cdc.gov/sth/about/ascariasis.html [Accessed JULY 20 2026].

Chehayeb, J. F., Robertson, A. P., Martin, R. J., Geary, T. G. 2014. Proteomic analysis of adult Ascaris suum fluid compartments and secretory products. PLoS Negl Trop Dis, 8, e2939.

Chen, J., Gong, Y., Chen, Q., Li, S., Zhou, Y. 2024. Global burden of soil-transmitted helminth infections, 1990-2021. Infect Dis Poverty, 13, 77.

Chow, F. W., Koutsovoulos, G., Ovando-Vázquez, C., Neophytou, K., Bermúdez-Barrientos, J. R., Laetsch, D. R., Robertson, E., Kumar, S., Claycomb, J. M., Blaxter, M., Abreu-Goodger, C., Buck, A. H. 2019. Secretion of an Argonaute protein by a parasitic nematode and the evolution of its siRNA guides. Nucleic Acids Res, 47, 3594–3606.

Clancy, J. W., Schmidtmann, M., D’souza-Schorey, C. 2021. The ins and outs of microvesicles. FASEB Bioadv, 3, 399–406.

Coronado, S., Barrios, L., Zakzuk, J., Regino, R., Ahumada, V., Franco, L., Ocampo, Y., Caraballo, L. 2017. A recombinant cystatin from Ascaris lumbricoides attenuates inflammation of DSS-induced colitis. Parasite Immunol, 39.

Coronado, S., Zakzuk, J., Regino, R., Ahumada, V., Benedetti, I., Angelina, A., Palomares, O., Caraballo, L. 2019. Cystatin Prevents Development of Allergic Airway Inflammation in a Mouse Model. Front Immunol, 10, 2280.

Cortés, A., Sánchez-López, C. M., Gónzalez-Arce, A., Bernal, D., Marcilla, A. 2025. Helminth extracellular vesicles: Roles in and beyond host-parasite communication. Adv Parasitol, 128, 1–34.

Cully, D. F., Vassilatis, D. K., Liu, K. K., Paress, P. S., Van der Ploeg, L. H., Schaeffer, J. M., Arena, J. P. 1994. Cloning of an avermectin-sensitive glutamate-gated chloride channel from Caenorhabditis elegans. Nature, 371, 707–11.

Cwiklinski, K., DE LA TORRE-Escudero, E., Trelis, M., Bernal, D., Dufresne, P. J., Brennan, G. P., O’neill, S., Tort, J., Paterson, S., Marcilla, A., Dalton, J. P., Robinson, M. W. 2015. The Extracellular Vesicles of the Helminth Pathogen, Fasciola hepatica: Biogenesis Pathways and Cargo Molecules Involved in Parasite Pathogenesis. Mol Cell Proteomics, 14, 3258–73.

De Silva, N. R., Brooker, S., Hotez, P. J., Montresor, A., Engels, D., Savioli, L. 2003. Soil-transmitted helminth infections: updating the global picture. Trends Parasitol, 19, 547–51.

Dennis, E. A., Cao, J., Hsu, Y. H., Magrioti, V., Kokotos, G. 2011. Phospholipase A2 enzymes: physical structure, biological function, disease implication, chemical inhibition, and therapeutic intervention. Chem Rev, 111, 6130–85.

Donnelly, S., O’neill, S. M., Sekiya, M., Mulcahy, G., Dalton, J. P. 2005. Thioredoxin peroxidase secreted by Fasciola hepatica induces the alternative activation of macrophages. Infect Immun, 73, 166–73.

Donnelly, S., Stack, C. M., O’neill, S. M., Sayed, A. A., Williams, D. L., Dalton, J. P. 2008. Helminth 2-Cys peroxiredoxin drives Th2 responses through a mechanism involving alternatively activated macrophages. FASEB J, 22, 4022–32.

Eichenberger, R. M., Sotillo, J., Loukas, A. 2018. Immunobiology of parasitic worm extracellular vesicles. Immunol Cell Biol.

EL Andaloussi, S., Mäger, I., Breakefield, X. O., Wood, M. J. 2013. Extracellular vesicles: biology and emerging therapeutic opportunities. Nat Rev Drug Discov, 12, 347–57.

Gao, X., Tyagi, R., Magrini, V., Ly, A., Jasmer, D. P., Mitreva, M. 2017. Compartmentalization of functions and predicted miRNA regulation among contiguous regions of the nematode intestine. RNA Biol, 14, 1335–1352.

Geary, T. G. 2005. Ivermectin 20 years on: maturation of a wonder drug. Trends Parasitol, 21, 530–2.

Hansen, E. P., Fromm, B., Andersen, S. D., Marcilla, A., Andersen, K. L., Borup, A., Williams, A. R., Jex, A. R., Gasser, R. B., Young, N. D., Hall, R. S., Stensballe, A., Ovchinnikov, V., Yan, Y., Fredholm, M., Thamsborg, S. M., Nejsum, P. 2019. Exploration of extracellular vesicles from ascaris suum provides evidence of parasite-host cross talk. J Extracell Vesicles, 8, 1578116.

Hansen, T. B., Jensen, T. I., Clausen, B. H., Bramsen, J. B., Finsen, B., Damgaard, C. K., Kjems, J. 2013. Natural RNA circles function as efficient microRNA sponges. Nature, 495, 384–8.

Harischandra, H., Yuan, W., Loghry, H. J., Zamanian, M., Kimber, M. J. 2018. Profiling extracellular vesicle release by the filarial nematode Brugia malayi reveals sex-specific differences in cargo and a sensitivity to ivermectin. PLoS Negl Trop Dis, 12, e0006438.

Harpur, R. P. 1977. Anatomy of the intestine of Ascaris suum: a three dimensional study with silicone rubber casts. Can J Zool, 55, 1110–7.

Hayden, M. S. C Ghosh, S. 2008. Shared principles in NF-kappaB signaling. Cell, 132, 344–62.

Hewitson, J. P., Harcus, Y. M., Curwen, R. S., Dowle, A. A., Atmadja, A. K., Ashton, P. D., Wilson, A., Maizels, R. M. 2008. The secretome of the filarial parasite, Brugia malayi: proteomic profile of adult excretory-secretory products. Mol Biochem Parasitol, 160, 8–21.

Janssen, I. J., Krücken, J., Demeler, J., Von Samson-Himmelstjerna, G. 2015. Transgenically expressed Parascaris P-glycoprotein-11 can modulate ivermectin susceptibility in Caenorhabditis elegans. Int J Parasitol Drugs Drug Resist, 5, 44–7.

Karabowicz, J., Długosz, E., Bąska, P., Wiśniewski, M. 2022. Nematode Orthologs of Macrophage Migration Inhibitory Factor (MIF) as Modulators of the Host Immune Response and Potential Therapeutic Targets. Pathogens, 11.

Kobpornchai, P., Reamtong, O., Phuphisut, O., Malaitong, P., Adisakwattana, P. 2022. Serine protease inhibitor derived from Trichinella spiralis (TsSERP) inhibits neutrophil elastase and impairs human neutrophil functions. Front Cell Infect Microbiol, 12, 919835.

Kubata, B. K., Duszenko, M., Martin, K. S., Urade, Y. 2007. Molecular basis for prostaglandin production in hosts and parasites. Trends Parasitol, 23, 325–31.

Kugeratski, F. G., Hodge, K., Lilla, S., Mcandrews, K. M., Zhou, X., Hwang, R. F., Zanivan, S., Kalluri, R. 2021. Quantitative proteomics identifies the core proteome of exosomes with syntenin-1 as the highest abundant protein and a putative universal biomarker. Nat Cell Biol, 23, 631–641.

Laing, R., Bartley, D. J., Morrison, A. A., Rezansoff, A., Martinelli, A., Laing, S. T., Gilleard, J. S. 2015. The cytochrome P450 family in the parasitic nematode Haemonchus contortus. Int J Parasitol, 45, 243–51.

Lamassiaude, N., Courtot, E., Corset, A., Charvet, C. L., Neveu, C. 2022. Pharmacological characterization of novel heteromeric GluCl subtypes from Caenorhabditis elegans and parasitic nematodes. Br J Pharmacol, 179, 1264–1279.

Leles, D., Gardner, S. L., Reinhard, K., Iñiguez, A., Araujo, A. 2012. Are Ascaris lumbricoides and Ascaris suum a single species? Parasit Vectors, 5, 42.

Lespine, A., Blancfuney, C., Prichard, R., Alberich, M. 2024. P-glycoproteins in anthelmintic safety, efficacy, and resistance. Trends Parasitol, 40, 896–913.

Liu, T., Zhang, L., Joo, D., Sun, S. C. 2017. NF-κB signaling in inflammation. Signal Transduct Target Ther, 2, 17023-.

Locksley, R. M. 1994. Th2 cells: help for helminths. J Exp Med, 179, 1405–7.

Loghry, H. J., Sondjaja, N. A., Minkler, S. J., Kimber, M. J. 2022. Secreted filarial nematode galectins modulate host immune cells. Front Immunol, 13, 952104.

Loghry, H. J., Yuan, W., Zamanian, M., Wheeler, N. J., Day, T. A., Kimber, M. J. 2020. Ivermectin inhibits extracellular vesicle secretion from parasitic nematodes. J Extracell Vesicles, 10, e12036.

Maizels, R. M., Mcsorley, H. J. 2016. Regulation of the host immune system by helminth parasites. J Allergy Clin Immunol, 138, 666–675.

Maizels, R. M., Yazdanbakhsh, M. 2003. Immune regulation by helminth parasites: cellular and molecular mechanisms. Nat Rev Immunol, 3, 733–44.

Manikantan, V., Ripley, N. E., Nielsen, M. K., Dangoudoubiyam, S. 2024. Protein profile of extracellular vesicles derived from adult Parascaris spp. Parasit Vectors, 17, 426.

Marcilla, A., Trelis, M., Cortés, A., Sotillo, J., Cantalapiedra, F., Minguez, M. T., Valero, M. L., Sánchez del Pino, M. M., Muñoz-Antoli, C., Toledo, R., Bernal, D. 2012. Extracellular vesicles from parasitic helminths contain specific excretory/secretory proteins and are internalized in intestinal host cells. PLoS One, 7, e45974.

Martin, R. J. 1996. An electrophysiological preparation of Ascaris suum pharyngeal muscle reveals a glutamate-gated chloride channel sensitive to the avermectin analogue, milbemycin D. Parasitology, 112 (Pt 2), 247–52.

Martin, R. J., Kusel, J. R., Pennington, A. J. 1990. Surface properties of membrane vesicles prepared from muscle cells of Ascaris suum. J Parasitol, 76, 340–8.

Martin, R. J., Robertson, A. P., Bjorn, H. 1997. Target sites of anthelmintics. Parasitology, 114 Suppl, S111–24.

Matoušková, P., Lecová, L., Laing, R., Dimunová, D., Vogel, H., Raisová Stuchlíková, L., Nguyen, L. T., Kellerová, P., Vokřál, I., Lamka, J., Szotáková, B., Várady, M., Skálová, L. 2018. UDP-glycosyltransferase family in Haemonchus contortus: Phylogenetic analysis, constitutive expression, sex-differences and resistance-related differences. Int J Parasitol Drugs Drug Resist, 8, 420–429.

Matoušková, P., Vokřál, I., Lamka, J., Skálová, L. 2016. The Role of Xenobiotic-Metabolizing Enzymes in Anthelmintic Deactivation and Resistance in Helminths. Trends Parasitol, 32, 481–491.

Memczak, S., Jens, M., Elefsinioti, A., Torti, F., Krueger, J., Rybak, A., Maier, L., Mackowiak, S. D., Gregersen, L. H., Munschauer, M., Loewer, A., Ziebold, U., Landthaler, M., Kocks, C., LE Noble, F., Rajewsky, N. 2013. Circular RNAs are a large class of animal RNAs with regulatory potency. Nature, 495, 333–8.

Minkler, S. J., Loghry-Jansen, H. J., Sondjaja, N. A., Kimber, M. J. 2022. Expression and Secretion of Circular RNAs in the Parasitic Nematode,. Front Genet, 13, 884052.

Moreno, Y., Nabhan, J. F., Solomon, J., Mackenzie, C. D., Geary, T. G. 2010. Ivermectin disrupts the function of the excretory-secretory apparatus in microfilariae of Brugia malayi. Proc Natl Acad Sci U S A, 107, 20120–5.

Omura, S., Crump, A. 2004. The life and times of ivermectin - a success story. Nat Rev Microbiol, 2, 984–9.

Pérez-Morales, D., Espinoza, B. 2015. The role of small heat shock proteins in parasites. Cell Stress Chaperones, 20, 767–80.

Ricciotti, E., Fitzgerald, G. A. 2011. Prostaglandins and inflammation. Arterioscler Thromb Vasc Biol, 31, 986–1000.

Roepstorff, A., Mejer, H., Nejsum, P., Thamsborg, S. M. 2011. Helminth parasites in pigs: new challenges in pig production and current research highlights. Vet Parasitol, 180, 72–81.

Rontogianni, S., Synadaki, E., Li, B., Liefaard, M. C., Lips, E. H., Wesseling, J., Wu, W., Altelaar, M. 2019. Proteomic profiling of extracellular vesicles allows for human breast cancer subtyping. Commun Biol, 2, 325.

Savina, A., Furlán, M., Vidal, M., Colombo, M. I. 2003. Exosome release is regulated by a calcium-dependent mechanism in K562 cells. J Biol Chem, 278, 20083–90.

Schnoeller, C., Rausch, S., Pillai, S., Avagyan, A., Wittig, B. M., Loddenkemper, C., Hamann, A., Hamelmann, E., Lucius, R., Hartmann, S. 2008. A helminth immunomodulator reduces allergic and inflammatory responses by induction of IL-10-producing macrophages. J Immunol, 180, 4265–72.

Thamsborg, S. M., Nejsum, P., Mejer, H. 2013. Impact of Ascaris suum in livestock . <em data-end=“15S7” data-start=“1564“>Ascaris: The Neglected Parasite, Elsevier.

Tritten, L., Ballesteros, C., Beech, R., Geary, T. G., Moreno, Y. 2021. Mining nematode protein secretomes to explain lifestyle and host specificity. PLoS Negl Trop Dis, 15, e0009828.

Tritten, L., Geary, T. G. 2018. Helminth extracellular vesicles in host-parasite interactions. Curr Opin Microbiol, 46, 73–79.

Van Niel, G., D’angelo, G., Raposo, G. 2018. Shedding light on the cell biology of extracellular vesicles. Nat Rev Mol Cell Biol, 19, 213–228.

Wang, T., VAN Steendam, K., Dhaenens, M., Vlaminck, J., Deforce, D., Jex, A. R., Gasser, R. B., Geldhof, P. 2013. Proteomic analysis of the excretory-secretory products from larval stages of Ascaris suum reveals high abundance of glycosyl hydrolases. PLoS Negl Trop Dis, 7, e2467.

Wang, Y., Lu, M., Wang, S., Ehsan, M., Yan, R., Song, X., Xu, L., Li, X. 2017. Characterization of a secreted macrophage migration inhibitory factor homologue of the parasitic nematode Haemonchus Contortus acting at the parasite-host cell interface. Oncotarget, 8, 40052–40064.

Welsh, J. A., Goberdhan, D. C. I., O’driscoll, L., Buzas, E. I., Blenkiron, C., Bussolati, B., Cai, H., DI Vizio, D., Driedonks, T. A. P., Erdbrügger, U., FALCON-Perez, J. M., Fu, Q. L., Hill, A. F., Lenassi, M., Lim, S. K., Mahoney, M. G., Mohanty, S., Möller, A., Nieuwland, R., Ochiya, T., Sahoo, S., Torrecilhas, A. C., Zheng, L., Zijlstra, A., Abuelreich, S., Bagabas, R., Bergese, P., Bridges, E. M., Brucale, M., Burger, D., Carney, R. P., Cocucci, E., Crescitelli, R., Hanser, E., Harris, A. L., Haughey, N. J., Hendrix, A., Ivanov, A. R., JOVANOVIC-Talisman, T., Kruh-Garcia, N. A., Ku’ulei-Lyn Faustino, V., Kyburz, D., Lässer, C., Lennon, K. M., Lötvall, J., Maddox, A. L., MARTENS-Uzunova, E. S., Mizenko, R. R., Newman, L. A., Ridolfi, A., Rohde, E., Rojalin, T., Rowland, A., Saftics, A., Sandau, U. S., Saugstad, J. A., Shekari, F., Swift, S., TER-Ovanesyan, D., Tosar, J. P., Useckaite, Z., Valle, F., Varga, Z., Van DER Pol, E., Van Herwijnen, M. J. C., Wauben, M. H. M., Wehman, A. M., Williams, S., Zendrini, A., Zimmerman, A. J., Théry, C., Witwer, K. W. C Consortium, M. 2024. Minimal information for studies of extracellular vesicles (MISEV2023): From basic to advanced approaches. J Extracell Vesicles, 13, e12404.

WHO. 2023. *Soil-transmitted helminth infections* [Online]. World Health Organization. Available: https://www.who.int/news-room/fact-sheets/detail/soil-transmitted-helminth-infections [Accessed].

Wiśniewski, J. R., Zougman, A., Nagaraj, N. C Mann, M. 2009. Universal sample preparation method for proteome analysis. Nat Methods, 6, 359–62.

Wolstenholme, A. J. 2012. Glutamate-gated chloride channels. J Biol Chem, 287, 40232–8.

Younis, A. E., Geisinger, F., Ajonina-Ekoti, I., Soblik, H., Steen, H., Mitreva, M., Erttmann, K. D., Perbandt, M., Liebau, E. C Brattig, N. W. 2011. Stage-specific excretory-secretory small heat shock proteins from the parasitic nematode Strongyloides ratti--putative links to host’s intestinal mucosal defense system. FEBS J, 278, 3319–36.

Zamanian, M., Fraser, L. M., Agbedanu, P. N., Harischandra, H., Moorhead, A. R., Day, T. A., Bartholomay, L. C., Kimber, M. J. 2015. Release of Small RNA-containing Exosome-like Vesicles from the Human Filarial Parasite Brugia malayi. PLoS Negl Trop Dis, 9, e0004069.

