## Supplementary material for "Inhibition of release of intestinal extracellular vesicles in *Ascaris suum* and immune modulation by the anthelmintic ivermectin": Supplemenatary Table 1

| Primer | Sequence |
|---|---|
| *Asu-glc-2* F | TGACGGACACCTGATGAACG |
| *Asu-glc-2* R | GTGCCTTGAAGCGTTGACAG |
| *Asu-glc-3* F | ACTCAACCACCTCCTACAACACA |
| *Asu-glc-3* R | TGAAGATTAGGAAGCAAGCCG |
| *Asu-glc-3* 1F | CTTACCTCGCGATAATAAGCCTG |
| *Asu-glc-3 1R* | AATGAGACCCACGAAACAATGA |
| *Asu-glc-3 2R* | TCCTCTGTTTCACCCAAACTTC |
| *Asu-glc-4 F* | TCCAGAATGAACGCAACGG |
| *Asu-glc-4 R* | ATCGCTTGTCGTATACGCGT |
| *Asu-avr-14 F* | ATGTTCCGAGTACAGAATGATT |
| *Asu-avr-14 R* | TCACATCAAGTAGACGGC |
| *Asu-avr-14 FLF* | ATGTTCCGAGTACAGAATGATT |
| *Asu-avr-14 FLR* | TCACATCAAGTAGACGGC |
| *Asu-gapdh F* | TCTCGAATGCATCCTGCACC |
| *Asu-gapdh R* | CACGTCCATCTCTCCATTGC |

**Supplementary Table 1: Primer sequences used for RT-PCR**. Forward (F) and reverse (R) primers targeting the glutamate-gated chloride channel subunit genes *Asu-glc-2*, *Asu-glc-3*, *Asu-glc-4*, and *Asu-avr-14*, with the housekeeping reference gene *gapdh* for positive control. Full-length *avr-14* primers used for Sanger sequencing are denoted FLF (forward) and FLR (reverse).
