## Supplementary Table 2 for "Inhibition of release of intestinal extracellular vesicles in *Ascaris suum* and immune modulation by the anthelmintic ivermectin"

| **Gene Name** | **Gene ID** | **Accession number** |
|---|---|---|
| *Asu-glc-2* | AgR035X_g084 | AEUI03000050.1 |
| *Asu-glc-3* | AgB19_g075 | AEUI03000059.1 |
| *Asu-glc-4* | AgB03_g128 | AEUI03000005.1 |
| *Asu-avr-14* | AgB04_g114 | AEUI03000006.1 |

**Supplementary Table 2: Gene identifiers and accession numbers for genes targeted by RT-PCR** WormBase ParaSite gene IDs and corresponding accession number for each of the four glutamate-gated chloride channel subunit genes (*Asu-glc-2*, *Asu-glc-3*, *Asu-glc-4*, and *Asu-avr-14*) targeted by the primer sets in Supplementary Table 1 from the *A. suum* genome PRJNA62057.
