## Supplementary Table 3 for "Inhibition of release of intestinal extracellular vesicles in *Ascaris suum* and immune modulation by the anthelmintic ivermectin"

| UniProt Accession | Identified Protein name | Protein function | Ivermectin Response & Expression Trend |
|---|---|---|---|
| F1KWT2 | Nuclear factor NF-κB p105 subunit | unknown | Down trend, significant. |
| F1LFX1 | Phospholipase A2 (PLA2) | *Steinernema carpocapsae*, whose secreted PLA2 suppresses host cellular and humoral immunity (downregulating Toll/Imd-pathway antimicrobial peptides and phagocytosis) in a *Drosophila* infection model (Parks et al., 2023). | Down trend, significant. |
| F1KZP5 | Small heat shock protein (sHSP) OV25-2 | Small heat shock protein (sHSP) family member identified among EV proteome heat shock proteins (Hansen et al., 2019)<br><br>*Brugia malayi*: its secreted sHSP BmHSP12.6 binds human IL-10 receptor alpha, blocks native IL-10 binding, and mimics IL-10-like activity via its N-terminal domain, and HSP12.6 orthologs are confirmed present in the *A. suum* genome, though never functionally tested(Gnanasekar et al., 2008, Dakshinamoorthy et al., 2012). | Down trend, significant. |
| F1LCK6 / F1LCD7 | Transthyretin-like (TTL) protein 46 / 5 | TTL family proteins are among the parasitism-associated protein families identified in *A. suum* larval ES products, expanded relative to free-living nematodes(Wang et al., 2013).<br><br>*Haemonchus contortus*: the secreted ortholog TTR-31 is required for germ-cell apoptotic clearance and post-embryonic larval development (Shi et al., 2021).<br><br>*Ostertagia ostertagi*: a broader survey across strongylid ES products found TTL family expansion with a hypothesized nervous-system signalling role (Saverwyns et al., 2008). | Down trend, significant. |
| F1KQN0 | CD109 antigen | CD109 antigen is an automated homology label for α2-macroglobulin (α2M) family members. Identifies α2-macroglobulin as an opsonizing protease inhibitor in invertebrate innate immunity that neutralizes exogenous (including parasite-secreted) proteases by tagging them for endocytosis and intracellular degradation, rather than by direct active-site inhibition (Armstrong, 2010). | Up trend, significant. |
| F1KPM2 | Apolipophorin | unknown | Down trend, significant. |
| F1KUF9 | ABA-1 / nematode polyprotein allergen (NPA) | ABA-1 is a lipid-binding nematode polyprotein allergen (NPA) with tandemly repeating ~15 kDa fatty-acid/retinol-binding subunits and stage/tissue-specific expression (Xia et al., 2000).<br><br>It is the dominant IgE target in human ascariasis, with IgE reactivity tracking host resistance/susceptibility to infection (McSharry et al., 1999). | Down trend, not significant. |

| | | | |
|---|---|---|---|
| | | Purified ABA-1 was found dispensable for the Th2-skewing immunomodulatory activity of *Ascaris* body fluid, which is instead driven by other body-fluid components (Paterson et al., 2002). | |
| F1LEI7 | FAR-1 (fatty-acid- and retinol-binding protein 1) | Recombinant FAR-1/FAR-2 from the insect-parasitic nematode *Steinernema carpocapsae* suppress the phenoloxidase cascade and antimicrobial peptide production and increase host susceptibility to bacterial co-infection in vivo, with similar immunosuppressive effects shown for FARs from *Ancylostoma ceylanicum* (Parks et al., 2021). | No detectable IVM effect. |
| F1KVP6 / F1LHQ3 | Cystatin | *Ascaris lumbricoides* cystatin (Al-CPI) prevents allergic airway inflammation and attenuates DSS-induced colitis in mouse models and upregulates the mevalonate/cholesterol biosynthesis pathway and immunomodulatory genes in human monocyte-derived dendritic cells (Coronado et al., 2017, Coronado et al., 2019, Acevedo et al., 2024).<br><br>A related filarial nematode cystatin drives IL-10-producing regulatory macrophages that reduce allergic/inflammatory responses (Schnoeller et al., 2008). | Down trend, not significant. |
| F1KV03 / F1LFD6 | Macrophage migration inhibitory factor (MIF) / MIF-like protein mif-2 | Nematode MIF orthologs directly bind host monocytes/macrophages and modulate their immunomodulatory activity, as shown for a secreted MIF homolog of the gastrointestinal nematode *Haemonchus contortus* acting at the parasite-host cell interface (Wang et al., 2017).<br><br>Nematode MIF orthologs more broadly are recognized as structural and functional mimics of host MIF with the potential to modulate the host immune response across multiple parasitic species (Karabowicz et al., 2022).<br><br>Host-derived MIF is itself required for effective Type 2 (Th2) effector immunity against the intestinal helminth *Heligmosomoides polygyrus* (Filbey et al., 2019).<br><br>MIF-deficient mice mount an enhanced Th2 response and clear the nematode *Nippostrongylus brasiliensis* more effectively than wild-type mice, showing that the direction of MIF's effect on anti-helminth immunity is host/species-context dependent (Damle et al., 2017). | No detectable IVM effect. |
| F1LAD2, F1L893, F1KZZ8, F1L9M1, F1LFS8 | Galectins | Galectins were detected among the excretory-secretory proteins of adult *A. suum*.(Midha et al., 2018).<br><br>Filarial nematode *Brugia malayi*, the EV-associated galectin Bma-LEC-2 is bioactive and polarizes host macrophages toward an alternatively activated (M2) phenotype, selectively induces apoptosis in Th1 cells, and increases macrophage IL-10 production (Loghry et al., 2022). | Up trend in IVM: F1L9M1, F1LAD2, F1LFS8. not significant<br>Down trend in IVM: F1KZZ8, F1L893. not significant |

| | | | |
|---|---|---|---|
| F1L4J8 | Serpin (TsSERPs) | Secreted nematode serpins directly inhibit host neutrophil elastase and cathepsin G, blocking neutrophil extracellular trap (NET) formation and phagocytic function, as shown for *Trichinella spiralis* TsSERP1(Kobpornchai et al., 2022).<br><br>A distinct serpin secreted by *Brugia malayi* microfilariae directly inhibits human neutrophil serine proteinases(Zang et al., 1999).<br><br>A recombinant serpin from *Trichinella pseudospiralis* drives macrophage polarization toward the M2 (alternatively activated) phenotype via activation of the JAK2/STAT3 signaling pathway (Xu et al., 2017). | Up trend in IVM. not significant |

**Supplementary Table 3. Putative immunomodulatory proteins identified in the *Ascaris suum* intestinal EV proteome.**

Functional evidence for potential immune-related proteins either directly from *Ascaris*, characterized homologs in filarial and/or trichinellid nematodes. The corresponding supporting literature is provided.
